# Predicting the unpredictable: heatwaves and history of variability shape phytoplankton community thermal responses within one generation

**DOI:** 10.1101/2022.07.20.500758

**Authors:** Maria Elisabetta Santelia, Luisa Listmann, Stefanie Schnell, Elisa Schaum

## Abstract

Predicting the effect of increased thermal unpredictability, for example in the shape of heatwaves on phytoplankton metabolic responses is ripe with challenges. While single genotypes in laboratory environments will respond to environmental fluctuations in predictable and repeatable ways, it is difficult to relate rapid evolutionary responses of whole communities to their ecological history. Previously experienced environments, including fluctuations therein, can shape an organism’s specific niche as well as their responses to further environmental changes. This is a testable hypothesis as long as samples can be obtained where the environmental history is known, sufficiently diverse, and not obscured by confounding parameters such as day length and precipitation patterns. Here, we tested immediate (i.e. within one generation) metabolic temperature responses of natural phytoplankton assemblages from two thermally distinct regions in the Baltic Sea: the warmer and less predictable Kiel Area, and the overall colder and more predictable Bornholm Basin. Our approach allows us to investigate effects on immediate physiological time scales (responses curves), ecological and evolutionary processes on longer time scales (seasonal differences between basins) as well as mid-term responses during a natural occurring heatwave. We found evidence for a higher degree of phenotypic plasticity in samples from unpredictable environments (Kiel Area).

## Introduction

Rising temperatures, more frequent heatwaves and increased unpredictability in sea surface temperature all have the potential to change the way phytoplankton communities contribute to whether coastal waters act as carbon sources, or carbon sinks. Here, we investigate how immediate (i.e. within one generation) metabolic responses in phytoplankton communities’ (whole community and pico-phytoplankton fraction) are shaped by a history of biotic relationships and physical conditions.

Warming strongly affects phytoplankton populations: for example, geographical distribution of phytoplankton communities may shift (Thomas *et al*, 2012) in addition to changes in community composition, e.g. higher occurrence of cyanobacteria blooms (Visser *et al*., 2016)and altered phenotypic trait expression. Photosynthesis (P) and respiration (R) in particular, are labile phenotypic traits, i.e. they exhibit high phenotypic plasticity (Schaum *et al*., 2017), where traits can change without there being underlying heritable changes to the genome. Phytoplankton are crucial components of the carbon cycle and determine whether surface waters act primarily as CO_2_ sources or sinks (Field *et al*, 1998). Therefore, it is crucial that we understand how rapidly photosynthesis and respiration can change in response to the projected temperature variations of the next decades. Metabolic rates dictate the amount of carbon that can be allocated for growth and basal cellular maintenance (Brown *et al*, 2004) and shape other responses related to fitness (e.g. mortality, abundance (e.g. Dell *et al*, 2011)), and competitive ability (Bestion *et al*., 2018). Numerous short-term studies over one or a few generations and usually carried out on single species have shown that respiration is overall more sensitive to warming than photosynthesis (López-Urrutia *et al*., 2006; Regaudie-de-Gioux *et al*., 2014; Laufkotter *et al*., 2015; Barton *et al*., 2020). Therefore, the amount of carbon available to cells for growth and other processes declines as temperature rises. On the ecosystem level, models predict that this may lead to a sharp decline in primary productivity (up to 20%) in the next decades (Boyce *et al*, 2010; Bopp *et al*., 2013). However, for more accurate predictions, models need multi-species data on evolutionary time scales.

The effect of temperature on traits is described by thermal reaction norms (Kingsolver, 2009). In ectotherms, they show a common pattern (Eppley, 1972; Kingsolver, 2009): rates increase exponentially with rising temperature up to an optimum, after which they decline abruptly. On a cellular level, this pattern is mainly driven by enzymatic constraints (Schoolfield *et al*, 1980; Lee and Peterson, 2008). On an ecological level, the shape of thermal reaction norms is driven by thermal tolerance and the interplay of acclimation and local adaptation (Padfield *et al*., 2016).

Thermal optima (T_opt_) are usually correlated with environmental mean temperature (Pawar *et al*, 2015; Schaum *et al*., 2017). This relationship determines the degree of local adaptation (Mitchell & Lampert, 2000; Souther & Graw, 2011), while other parameters describing the shape of thermal reaction norms give indication about thermal tolerance. It describes the range of temperatures at which an organism or a population can survive, and can change and adapt as the environment changes (e.g. Bennett & Lenski, 2007; Lande, 2014; Thomas *et al*, 2016). Information on temperatures that phytoplankton communities can tolerate is crucial for a better understanding about the extent to which reaction norms of a trait can stay the same across environments (Magozzi & Calosi, 2015; Pacifici *et al*., 2015).

Sea surface temperature is naturally variable and largely depends on geographical and seasonal pattern (Karl *et al*., 2003). In the future, unpredictability is expected to increase, with a rise in frequency and severity of anomalous events such as heatwaves (Karl and Trenberth, 2003). Theory predicts that frequency and predictability of the environmental fluctuations will have an impact on the shapes of thermal tolerance curves in the short-term and long-term and thereby determines which strategies allow populations to persist in the environment. This can range from phenotypic plasticity to bet hedging (Botero *et al*., 2015). Ultimately, either strategy serves to avoid extinction and to increase the variance and mean in fitness (Starrfelt & Kokko, 2012).

Theoretical predictions are based on models and yet to be thoroughly tested in ecologically relevant scenarios. In natural populations, species persistence and adaptive trajectories are already shaped by the physical conditions they have faced, e.g. within species phenotypic variation is often high and can be partially explained by the environmental variability at the sampling location (Godhe & Rynearson, 2017). Theory and observational studies agree that local adaptation contributes to shaping adaptive and plastic trajectories organisms would potentially follow if exposed to new conditions (e.g. Pörtner, 2002; Boyd *et al*., 2016; Thomas *et al*, 2016). For phytoplankton, a strong correlation between mean temperature of the environment and thermal niches has been clearly demonstrated, showing a general large scale pattern of local adaptation, with tropical species tending to have thermal optima closer to the mean experienced temperature, probably due to thermal constraints at higher temperatures (Thomas *et al*., 2012).

We use the term eco-evolutionary history to describe the combination of biotic and abiotic factors that an organism has previously experienced and that have likely influenced the organisms’ adaptive trajectories. Specifically, past eco-evolutionary history can either influence the speed at which organisms adapt to novel and changing environments (Andrade-Restrepo *et al*, 2019) or, if ecosystems change on the same time scale as evolution occurs, it can be translated into changing community composition or interaction between species (Post & Palkovacs, 2009; Padfield *et al*., 2017).

Upscaling single species’ responses to the population level may not be a good predictor for population responses (Wolf *et al*, 2018). Community responses are often more than the sum of their parts and whole assemblages may respond in different ways to environmental parameters: trait-expression on the community level may stay unaltered e.g. via shifts at species or functional group level, or when individuals show a high degree of phenotypic plasticity (Godhe & Rynearson, 2017; Hoppe *et al*., 2018). Shifts in community composition are one of the main consequences of climate change. Evidence suggests warming is likely to increase the relative proportion of pico-phytoplankton, the smallest fraction of aquatic autotrophs (Morán *et al*., 2010). This shift toward environments dominated by smaller cells could potentially disrupt aquatic food webs and alter carbon export fluxes (Falkowski *et al*, 1998).

Understanding how the composition of communities and metabolic rates of phytoplankton change globally is a complicated task. Specifically, global scale studies face confounding factors, e.g. different light intensity and nutrient availability in spatially distant environments. Here, to minimise these confounding variables, we obtained natural phytoplankton communities from two well-characterised areas of the South-Western Baltic Sea: the Kiel Area and the Bornholm Basin. The two regions are close enough to be connected by currents, but at the same time, offer the rare chance to explore how a history of thermal variability may influence metabolic responses. Specifically, the Kiel Area is on average warmer and more thermally unpredictable than the more predictable and colder Bornholm Basin (Snoeijs-Leijonmalm & Andrén, 2017). We leveraged a series of cruises spanning two years, including a summer heatwave, to test how phytoplankton communities’ (whole community and pico-phytoplankton fraction) immediate metabolic responses to temperature are shaped by a history of biotic relationships and physical conditions. Responses within one generation - often before cells are fully acclimated to novel conditions – are important in order to avoid acclimations steps that can potentially alter the physiological responses of organisms to the considered variables(Munguia & Alenius, 2013).

## Materials and Methods

### Sampling areas

We chose two areas in the South-Western Baltic Sea, (Kiel/Mecklenburg Area and Bornholm Basin, abbreviated to KA and BB throughout) as sampling sites (Fig.1 A). We consider areas corresponding geographically to the Kiel Basin and the Mecklenburger Bight to belong to the same region (KA, Kiel Area) as we found them to be statistically indiscernible in their physical and chemical (e.g. salinities, nutrient concentrations) characteristics (see table S1).

**Figure 1.**
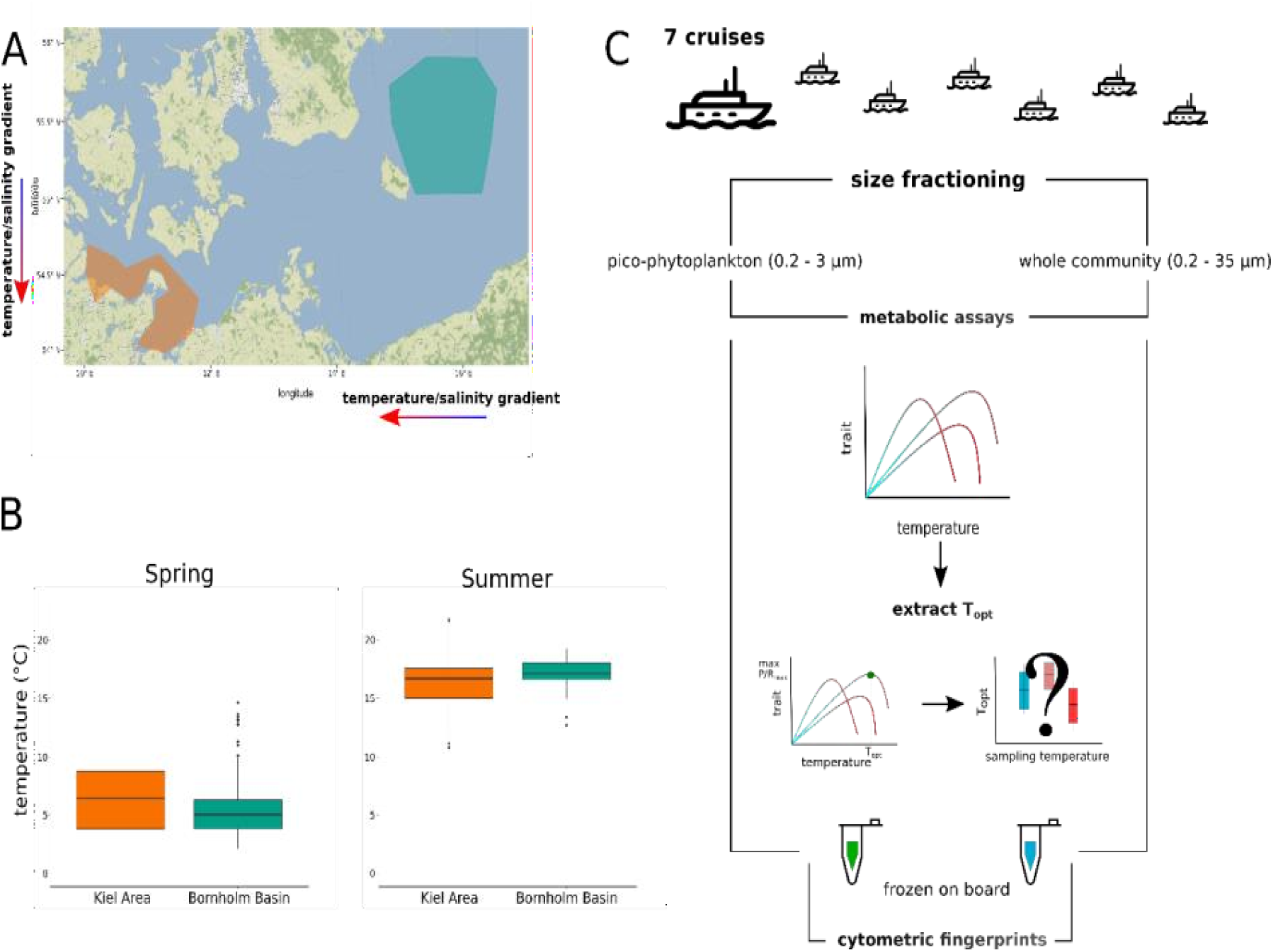
Sampling sites and experimental protocol flow chart. (**A**) Color-coded polygons describe the sampling areas (orange: Kiel Basin and Mecklenburger Bight, blue: Bornholm Basin). Arrows show direction of naturally occurring temperature and salinity gradients with red for warmer temperature and higher salinities, blue for colder temperatures and lower salinities. (**B**) Boxplots of surface temperatures (∼ 8 m) of the last 5 years for the Kiel Area, in orange, and the Bornholm Basin, in blue, for spring and summer. As data indicate sea surface temperature (SST) to be significantly more variable in the Kiel Area than in the Bornholm Basin, these two regions were chosen as sampling areas. **(C)** Infographic describing the sampling and consequent analyses on board and in the lab.

Over a period of two years, we sampled during the following cruises on RV ALKOR: March 2018 (AL505); July 2018 (AL513); March 2019 (AL520); April 2019 (AL521); May 2019 (AL522); July 2019 (AL524); October 2019 (AL530).

We took samples from at least 3 stations from each area on each cruise.

### Abiotic environment

We determined abiotic environmental conditions via i) decomposition analysis and ii) water sampling and nutrient measurements during the cruises prior to sample collection. We carried out a decomposition analysis using the function decompose of the *anomalize* package (0.2.0) in R version 4.0.2 to test for the differences in trend, seasonality and random impacts of temperature for the two chosen areas. Both additive (seasonal + trend + random) and multiplicative (seasonal * trend * random) approaches were used to analyse the residuals and anomalies after accounting for trend and seasonality. The additive approach assumes a quarterly seasonality (frequency = 4), whereas the multiplicative approach assumes a monthly seasonality (frequency = 12). Data used for the decomposition analysis contained monitoring temperature data of the last 5 years collected during ALKOR and POSEIDON’s Oceanographic cruises (data courtesy of the “GEOMAR – Helmotz Center for Ocean Research” of Kiel and “BSH - Bundesamt für Seeschiffahrt und Hydrographie” Center).

Before the water sample collection, physical and biological parameters of the water column during the cruises (position, temperature, salinity, chlorophyll a), were determined with a CTD (Conductivity, Temperature, Depth Water Sampler) probe (Table 1). Water samples for measuring dissolved inorganic nutrient concentrations were collected at 5 metres (crane-controlled Niskin Bottle or CTD Rosette) for each cruise and station. Samples were passed through an 0.2µm pore size filter and immediately stored frozen at −20 °C for subsequent colorimetric determination of nitrate, nitrite, ammonia, silicate and soluble reactive phosphorous (SRP) using a segmented flow auto-analyser (SEAL Analytics AA3, UK), following the methods of Murphy & Riley (1962), Kirkwood (1996), Grasshoff *et al* (2009).

**Table 1.** Overview of water column parameters during the cruises. Geographical areas refer to the two considered regions (KA: Kiel and Mecklenburger Basin; BB: Bornholm Basin). Data were retrieved from a CTD run at each station at the sampling depth (5-10 m). Each station was sampled only once as the water column up to sampling depth was fully mixed. Data were then pooled by sampling area

| Cruise date | Geographical Area | Salinity (PSU) | Water Temperature (° C) |
| --- | --- | --- | --- |
| AL505 (March 2018) | KA | 9.5 ± 4.64 | 1.34 ± 0.199 |
| AL505 (March 2018) | BB | 7.432 ± 0.09 | 2.32 ± 0.293 |
| AL513 (July 2018) | KA | 14.44 ± 0.433 | 21.03 ± 2.39 |
| AL513 (July 2018) | BB | 7.242 ± 0.433 | 20.88 ± 4.74 |
| AL520 (March 2019) | KA | 15.571 ± 2.09 | 4 ± 0 |
| AL521 (April 2019) | KA | 14.1 ± 4.192 | 6.26 ± 0.336± |
| AL521 (April 2019) | BB | 8.04 ± 0.09 | 6.4 ± 0.308 |
| AL522 (May 2019) | KA | 14.375 ± 1.888 | 11.225 ± 0.519 |
| AL522 (May 2019) | BB | 7.98 ± 0.045 | 9.36 ± 0.434 |
| AL524 (July 2019) | KA | 12.667 ± 1.53 | 18.4 ± 0.693 |
| AL524 (July 2019) | BB | 7.183 ± 1.059 | 17.167 ± 0.314 |
| AL530 (October 2019) | KA | 16.82 ± 0.96 | 12.92 ± 0.521 |

### Biological Sample collection

Water samples for measuring metabolic responses were collected with a Multi-Niskin bottles rosette sampler. Sampling depth was adjusted according to CTD profiles to make sure samples were taken above the thermocline and halocline if necessary but remained within the first 5-10 metres of the water column. Water samples were immediately passed through a 35 µm mesh to remove large grazers and particles. We then prepared two size fractions: one, containing all phytoplankton smaller than 35 µm, and another one, containing only pico-phytoplankton smaller than 3 µm. The former was passed through the 35 µm mesh and then concentrated on a 0.2 µm filter to increase biomass. The latter, pico-phytoplankton community sample, was filtered through 3 µm filters (Worden & Not, 2008), and the filtrate then concentrated on a 0.2 µm filter. We used at least 2 litres of water for each size fraction. From here onwards we will refer to the concentrated water samples containing either the 0.2 – 3 µm sized community or 0.2 – 35 µm sized community as “samples”.

### Metabolic Activity

To assess whether the organisms in the samples from different regions and cruises differed in their immediate (i.e. within one generation) metabolic responses to warming, we measured the rates of oxygen evolution in the light (Photosynthesis, P) and oxygen consumption in the dark (Respiration, R), using a Clark-type electrode (Oxytherm, HansaTech, UK). Gross photosynthesis (GP) was calculated as P + |R|, accounting for oxygen consumption both in the light and in the dark. For further analysis, we only used data, where photosynthesis was more positive than respiration, since when R exceeds P, the system points toward heterotrophy as a source of CO_2_ (del Giorgio *et al*, 1997). We used an aliquot of each sample to measure a Photosynthesis - Irradiance (PI) curve, to identify the optimal irradiance (I_opt_) at which to carry out measurements across a temperature gradient. Based on research carried out on phytoplankton communities, we expected I_opt_ to not change during immediate response measurements (Schaum *et al*, 2017).

Photosynthesis was measured at light intensities that spanned from 50 to 1500 µmol m^-2^ s^-1^ (with unequal increments, see supporting information) over 20 minutes (one minute at each light intensity) and followed by a 3 minute measurement of respiration in the dark. PI curves were then analysed in the R environment (v 3.5.3) using a modified Eilers’ photoinhibition model (Eilers & Peeters, 1988) (eq. 1),

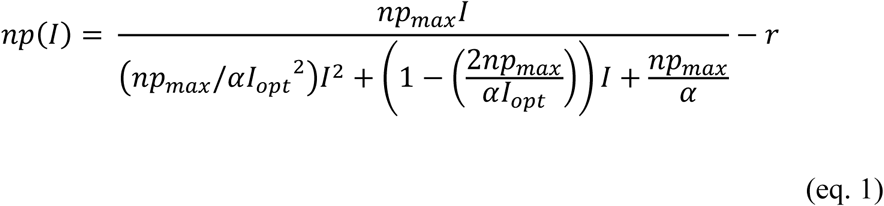

where *np*(*I*) is the rate of gross photosynthesis at the specific irradiance, *npmax* is the photosynthetic activity at the optimal light intensity (*Iopt*), α is the initial slope and *r* is the respiration when light intensity is 0.

Data were fitted using the package “*TeamPhytoplankton*”, based on non-linear least square regression, and the best fits were determined based on AIC scores running 1000 different combinations of initial parameters.

To ensure that the full breadth of the unimodal curve was captured, the range of assay temperatures for each cruise was based on trials carried out in the beginning of a cruise, as we did not expect parameters to drastically change over the sampling period. The temperature range included the lowest and the highest temperature at which a physiological response was reliably measurable. This resulted in ranges spanning from 3° up to 40° C in 1, 2 or 3° C increments (smaller increments near the thermal optimum, see Table S2 for details). No further measurements were carried out after temperatures at which oxygen consumption in the dark exceeded oxygen production in the light. Prior to the measurements, samples were given time to adjust to the assay temperature and conditions in the Oxytherm chamber until the respiration signal in the dark stabilised (max. 10 minutes). P and R were measured for 5 minutes each. A new aliquot was used for each assay temperature, to avoid stress responses or hysteresis induced by samples’ being subjected to multiple temperatures.

### Analysis of thermal reaction norms

Thermal reaction norms of metabolic rates of GP and R obtained on the oxygen electrode were subsequently analysed in the R environment (v 3.5.3), via non-linear least squares regression as stated above, using a modified Sharpe-Schoolfield equation (Schoolfield *et al*, 1980) (eq. 2),

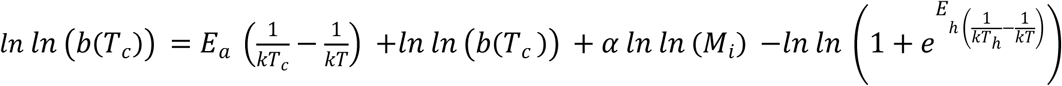

where *k* is the Boltzmann’s constant (8.62 × 10-5 eV/k), *E*_*a*_ is the activation energy (how *ln*(*b*(*T*_*c*_)) increases below the optimal temperature), *E*_*h*_ is the high-temperature induced inactivation of enzyme kinetics and *b*(*T*) the metabolic trait at the assay temperature (either GP or R). The optimum temperature was identified solving the following equation (eq. 3)

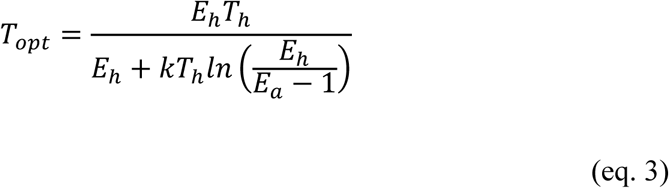

*E*_*a*_,*E*_*h*_ and *T*_*opt*_ represent biologically relevant features characterising thermal reaction norms. We extracted the T_opt_ values of GP and R for each samples and cruises thermal response curve and plotted them against *in situ* temperatures at the time of sampling.

### Flow cytometry

We collected a 200 µL aliquot of each sample processed at each tested temperature and froze it in sorbitol after measuring the metabolic rates (10 µL of a 1% sorbitol solution per sample). To ensure a gentle freezing process, samples were stored immediately at +4° C in the dark after adding sorbitol and then frozen at -80° C. Storage at +4° C depended on how fast samples could be processed on board, but never exceeded 48 hours. Cell numbers for biomass correction and phenotypic diversity were determined on a flow cytometer (Accuri C6, BD Biosciences, USA) upon returning to Hamburg. In order to avoid background noise and counting of heterotrophs or debris, we applied thresholds for both side scatter (measure of cellular size) and on the FL3 (chlorophyll) fluorescence channel.

The flow cytometry data were also used to establish flow cytometric fingerprints after Carr *et al* (2016). Even though analytical flow cytometry does not allow taxonomically relevant discrimination, the high functional (reflected in photopigment diversity) and morphological (both in terms of size and intracellular composition) diversity, allows distinction of phytoplankton clusters (for an overview of nomenclature used, see Table 2).

**Table 2.**
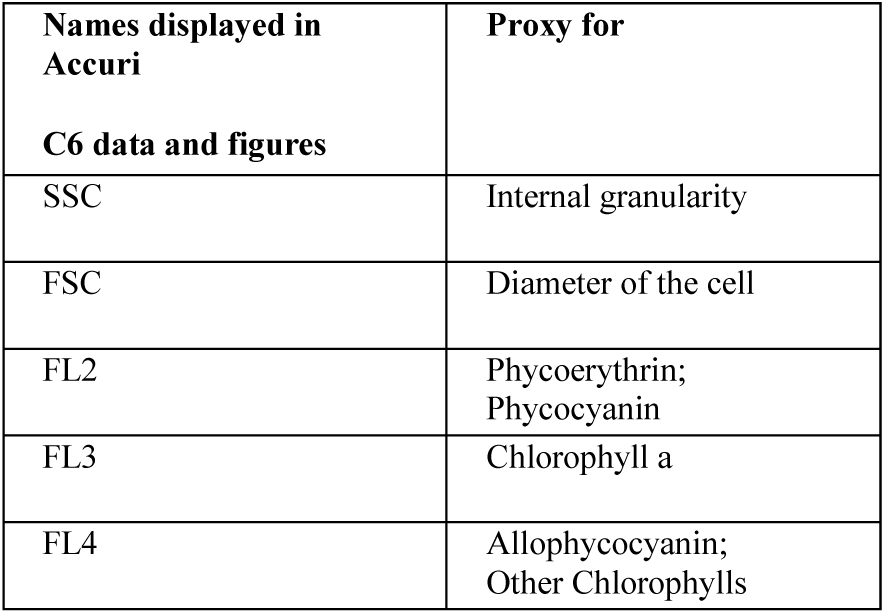
Nomenclature used for flow cytometry parameters used as proxies for phenotypic characteristics of the phytoplankton communities.

We also froze samples for bacteria quantification. In order to exclude geographically and/or seasonally driven differences in the bacteria concentration that could have shaped respiration patterns, we measured bacterial content of the aforementioned samples, upon returning to Hamburg. Prior to the cytometry analysis, samples were quickly thawed at 37° C and stained with SYBR Green I nucleic acid (Molecular Probes Inc., USA). Purchased dye has a 10000 fold concentration. SYBR Gold was diluted to a final concentration of 10^−4^ in TE buffer filtered on a 0.02 µm pore size filter prior to use, according to the method of Brussaard et al. (2000). Then the samples were first incubated at 80° C for 10 minutes and then cooled down in the dark before adding reference beads (1 µm diameter BD Biosciences, USA). Samples were then checked using a BD Accuri C6 flow cytometer, correcting for a blank with TE buffer and SYBR Green I prepared as the analysed samples. Bacteria concentration was established using scatter plots of green fluorescence of the staining (FL1 channel) versus side scatter as a measure of cells size (FSC channel).

### Data analysis and statistics

Data were processed in the R environment (version 4.0.2). Figures were made using *ggplot2* (3.2.1) and maps using *ggmap (3*.*0*.*0)*. We compared the extracted Topt values for GP and R using a mixed effects linear model using the package *nlme*. We fitted a global model including mean temperature at sampling time, geographical areas and size fraction as fixed and interacting (geo * fraction) effects and considered sampling stations as nested random effects. The model was subsequently simplified and models compared considering AIC scores and delta AIC values using the package *MuMIn* (1.43.6). In particular, we discarded models with Δ*i* >2. Δ*i* was calculated as the difference between *AICi* and *AICmin*, where *AICi* is the AIC score for the *i*th model and *AICmin* is the minimum of AIC among all models

Pairwise comparisons of slopes of linear regression between Topt and mean sampling temperature, were analysed examining the ANOVA p-values from interaction between mean sampling temperature, geographical areas and size fraction, then slopes were compared using the *lstrends* function. We used the *pairs* function for running a Tukey post-hoc comparison on the family of estimates. Both functions are built in the package *emmeans* (1.4.8).

To address whether samples from different cruises varied significantly in terms of microbial community composition, we conducted a Bayesian Principal Component Analysis (PCA) in the package *FactoMineR* (2.0) on the cytometric fingerprints (SSC, FSC, FL1, FL2; FL3; FL4). To test the differences of the community composition between the two geographical areas for both size fractions, a permutational Multivariate Analysis of Variance (PERMANOVA) was done using function *adonis* in the package *vegan* (2.5-6) using Bray-Curtis dissimilarity with 999 permutations on Euclidean matrix distances.

## Results

### Long-term monitoring data and sampling conditions

To chose sampling locations, we tested the difference in sea surface temperature variability of the two areas prior to the cruises and consequently chose sampling locations, and analysed environmental monitoring data. We separated spring and summer temperature data of the last 5 years (Fig. 1B). We found that the chosen areas (Kiel Area and Bornholm Basin) indeed showed different patterns in variability (Levene test: spring values, F_1,475_= 12.96, p=0.00035; summer values, F_1,383_= 4.21, p=0.04), with the Kiel Area’s standard deviation being overall higher than the standard deviation found in the Bornholm Basin (spring: Kiel Area ±3.36, Bornholm Basin ±2.86; summer: Kiel Area: ±2.07, Bornholm Basin: ±1.49). A decomposition analysis also confirmed the expected differences in predictability of temperature variations (Fig. S1): When decomposing the time series, the function was not able to predict a clear seasonal pattern for the Kiel Area. The random component was substantial for the Kiel Area as well, but this was not the case for data from the Bornholm Basin (One-Way ANOVA comparing random components outcomes comparing the two geographical areas: F_1, 2851_=128.9, p<2e-16; see Table S4 for statistics and random components).

The abiotic parameters (temperature, salinity, nutrients) measured during the cruises in the Kiel Area and Bornholm Basin followed a characteristic seasonal pattern: dissolved nitrogen, phosphate and silicate were replenished pre-bloom (March 2018/2019) (N: 73.92 µg L^-1^ ± 26.7; P: 16.12 ± 4.21 µg L^-1^; Si: 12.05 µM L^-1^ ± 1.14). As the spring and successive summer blooms established, all nutrient concentrations were reduced (July 2018/2019) (N: 14.96 µg L^- 1^ ± 4.45; P: 2.65 µg L^-1^ ± 0.51; Si: 4.54 µM L^-1^ ± 0.63) (Fig S2). There were no significant differences for temperature, or in major nutrient content, between the sampling regions during spring and summer (one-way ANOVA type III; temperature: F_1, 17_ = 0.02 p = 0.90; nitrate F_1,15_: 1.19, p = 0.29; phosphate F_1,15_: 0.11, p=0.74). Salinity was consistently lower in the Bornholm Basin than in the Kiel Area (Table 1; F_1,17_ = 24.87, p = 0.0001). There was a strong signal for abnormally long, abnormally warm periods during the summer cruise of 2018 (KA: 21.03 °C ± 2.18; BB: 20.88 °C ± 4.24) and to a smaller extend, during the summer cruise of 2019, while salinity and nutrient composition were comparable to previous years and long-term monitoring data (Snoeijs-Leijonmalm & Andrén, 2017). The summer of 2018 falls into the definition of heat wave given by Hobday *et al* (2016), with mean temperatures for both areas above the 90^th^ percentile for a period longer than 5 successive days.

### Mean *in situ* temperatures influence gross photosynthesis but not respiration thermal optima under average thermal conditions

In order to evaluate conditions reflecting the 90^th^ percentile of the past five years’ temperature average, we excluded the heat wave data from summer 2018 from our first round of analysis.

Under conditions excluding the heatwave, the shapes of acute thermal response curves of gross photosynthesis were highly malleable with regards to their optimum temperature (see Fig 2 A and B for thermal optima of gross photosynthesis (T_opt_GP) patterns over *in situ* temperatures). This reflects an ability of phytoplankton communities to alter the shape of the curve according to the environmental conditions on time scales of a single growing season, with higher T_opt_ (highest T_opt_ measured: 36.96 and 35.64 °C, respectively for Kiel Area and Bornholm Basin) at higher sea surface temperatures (highest sea surface temperature 21.03 and 20.09 °C for Kiel and Bornholm).

**Figure 2.**
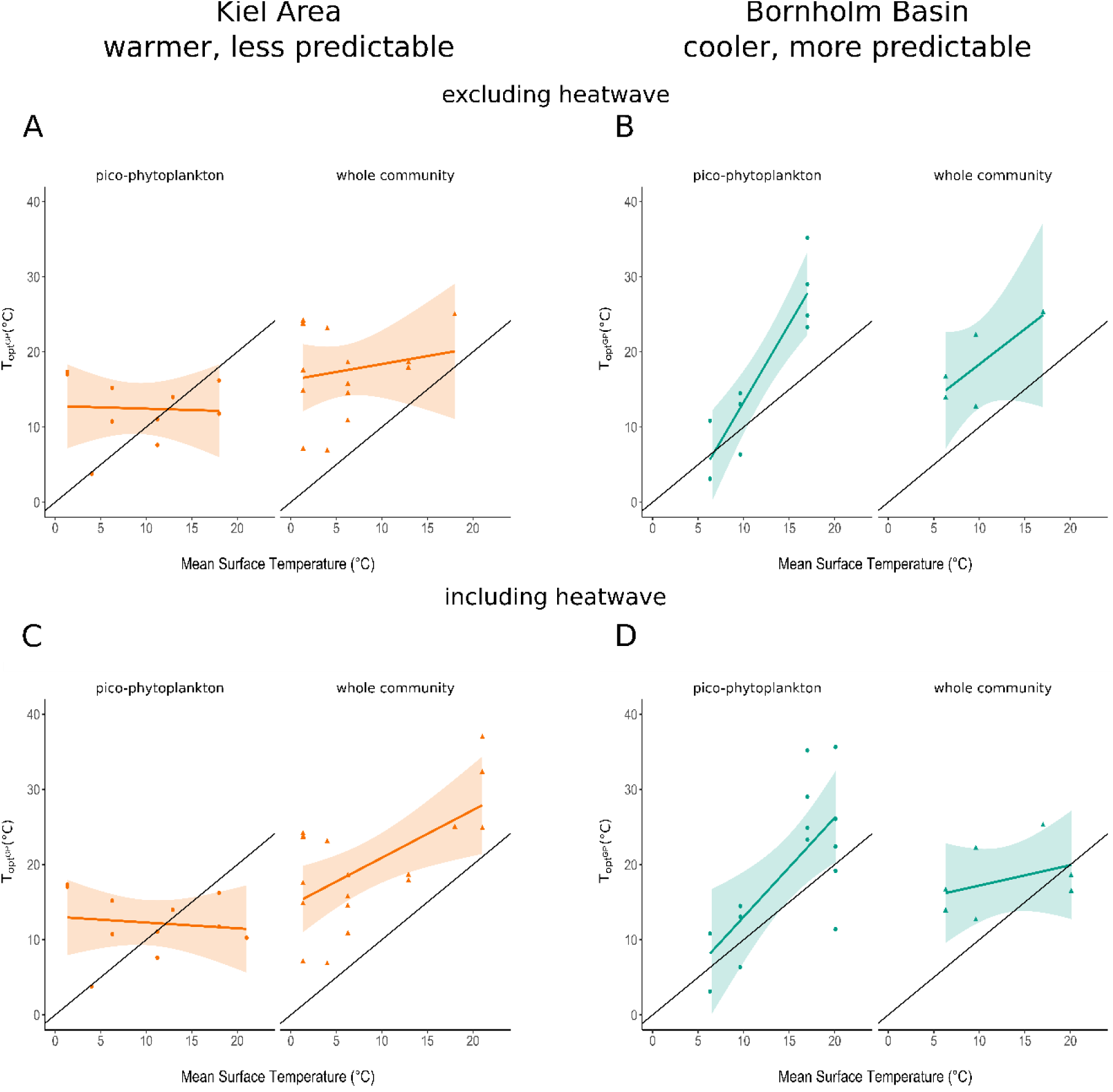
Scatter plot referring to the thermal optima of gross photosynthesis for the pico-phytoplankton and whole community in the two sampling areas (Kiel Area in orange and Bornholm Basin in blue). T_opt_GP was calculated according to eq. 3. The X-axis represents the mean surface water temperature at the time of the sampling in ºC. Coloured lines represent the output of linear regression with the corresponding confidence interval and the solid black line corresponds to the 1:1 regression between mean environmental temperature and thermal optima. Shaded areas are 95% confidence intervals automatically calculated in R. The first row (panel A and B) shows data excluding the 2018 summer heatwave. Second row (panel C and D) refers to the complete dataset, spanning the seven cruises of 2018 and 2019.

Specifically, samples from both size fractions from the Bornholm Basin and the whole community fraction from Kiel Area, showed an increase in thermal optima with increasing temperature whereas the pico-phytoplankton fraction from the Kiel Area showed a stable trend in thermal optima (Fig. 2 A and B). Thus, we found a significant effect of temperature and size fraction on the T_opt_GP especially regarding the whole phytoplankton community (mean sampling temperature: F_1,31_=5.65 p=0.024; size fraction: F_1,9_=11.64 p=0.008). In the pair-wise comparison of the slopes of the described trends between the different size fractions and geographical areas, we found that the pattern in the pico-phytoplankton Kiel Area fraction was significantly different from the other depicted trend for the pico-phytoplankton fraction in Bornholm Basin (Table 3).

**Table 3.** Pairwise comparisons of slopes of regression for T_opt_GP (thermal optima for gross photosynthesis) and mean sampling temperature, divided per geographical area of origin and size fraction. Only significant values are stated here. (SE: standard error; df: degree of freedom; p: p-value).

| Variable | SE | df | t.ratio | p | estimate |
| --- | --- | --- | --- | --- | --- |
| Excluding heatwave: pico-phytoplankton BB Vs. pico-phytoplankton KA | 0.399 | 38 | 3.810 | 0.0027 | 1.521 |
| Including heatwave: pico-phytoplankton BB Vs. pico-phytoplankton KA | 0.43 | 42 | 3.28 | 0.011 | 1.4 |

In contrast to the responses of GP, thermal optima for respiration (T_opt_RESP) did not change with increasing sea surface temperature in any of the considered areas and size spectra (Table S4 B for model comparison and ANOVA’s outputs for the most parsimonious model). Further, there were also no differences in the pairwise comparison of the slopes.

### Community respiration changes greatly during a heatwave

To evaluate whether the thermal optima substantially changed during a heatwave, we subsequently analysed the entire dataset including the extreme event of July 2018. Extreme events overall reinforced the relationship between T_opt_GP and seasonal changes in sea surface temperature (Fig. 2 C and D) (F_1,39_ = 13.83 p = 0.0006). Moreover, pairwise comparisons of the regression slopes additionally showed significant differences within the geographical areas and the size fraction. Specifically, we found differences between the pico-phytoplankton fraction from the two different Basins (Table 3). While inclusion of extreme events only had a small effect on responses of T_opt_GP, we found that T_opt_RESP was more responsive to extreme scenarios than during an average seasonal scenario (i.e. excluding the heatwave event), i.e. T_opt_RESP was strongly influenced by mean surface temperature (Fig. 3 C and D) (F_1,34_ = 21.97 p < 0.001). This response was conserved across all samples, regardless of size fraction or region of origin.

**Figure 3.**
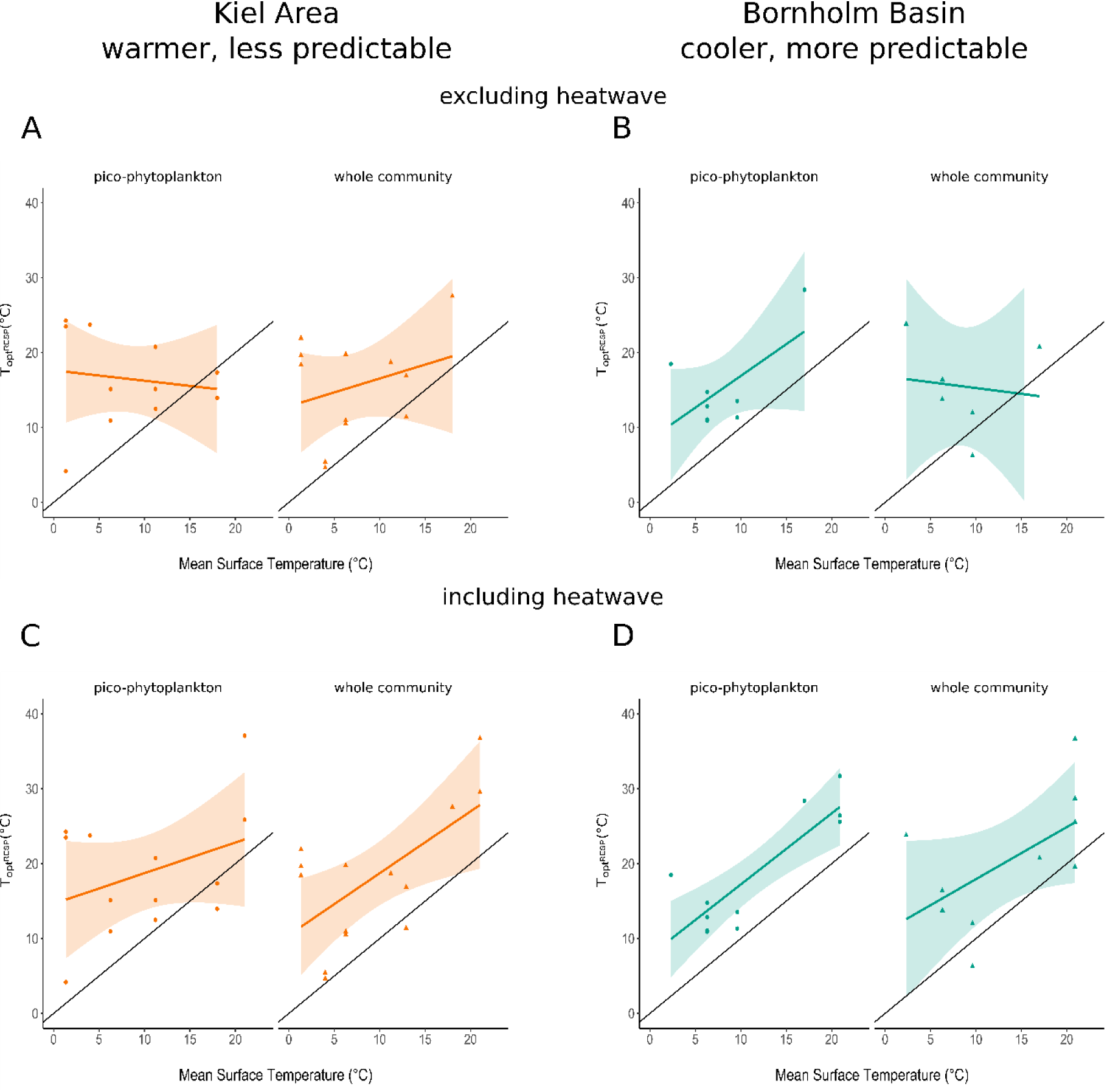
Scatter plot referring to the thermal optima of respiration (T_opt_RESP) for the pico-phytoplankton and whole community in the two different areas (Kiel Area in orange and Bornholm Basin in blue). The X-axis represents the mean surface water temperature at the time of the sampling in ºC. Coloured lines represent the output of linear regression with the corresponding confidence interval and solid black line corresponds to the 1:1 regression between mean environmental temperature and thermal optima. Shaded areas are 95% confidence intervals automatically calculated in R. The first row (panel A and B) shows data excluding the 2018 summer heatwave. Second row (panel C and D) refers to the complete dataset, spanning the seven cruises of 2018 and 2019.

### Changes in community composition offer an explanation for the changes of thermal optima responses in predictable regions but not for variable environments

In the absence of genetic community data (but see Zhong *et al* (2020) for a rough estimate of 2018 pico-phytoplankton community composition through meta-barcoding), we tracked the gross community composition on the functional group level throughout the cruises quantifying the covariance of phenotypic characteristics defined across cruises. In order to depict phenotypic features, we analysed the cytometric fingerprints, which describe the size and photopigments’ characteristics of the cells, for all the cruises and both size fractions. Each cruise is here throughout described as mean *in situ* temperature at the time of sampling to reflect environmental conditions.

We detected two trends: one, organisms sampled in the Kiel Area were phenotypically – at least in terms of size, granularity and photopigments – similar across seasons (and thereby temperatures). This is evidenced graphically by the clusterization of cytometric fingerprints of all the cruises (PERMANOVA; for the pico-phytoplankton and whole community, p>0.1; Fig. 4). Two, in the Bornholm Basin the clusters corresponding to the cruises were significantly different from each other, indicating that gross community composition changed (p<0.005, see also Table S5 for statistics; Fig. 5). In both size fractions and geographical areas, the main drivers of community distinction was not due to the changes in SSC (granularity) and FCS (size), but rather photopigment characteristics. The trend was preserved regardless of whether data obtained during the heatwave was included in the dataset (Fig 4 C and D; Fig. 5 C and D).

**Figure. 4.**
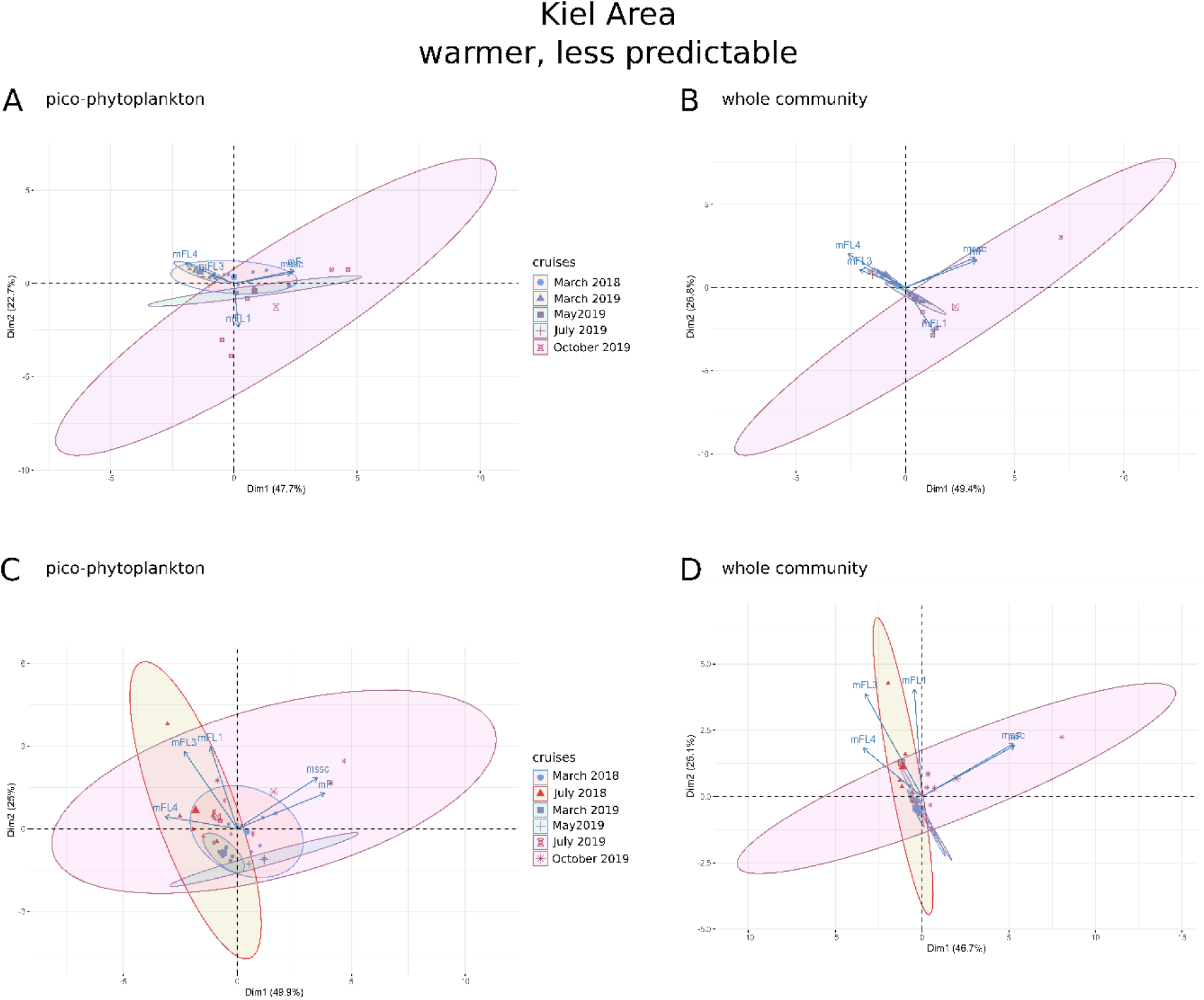
PCA of cytometric fingerprint covariance in the Kiel Area whole community and pico-phytoplankton fraction. Loadings represent phenotypic characteristics of functional groups derived from flow cytometry. The first row (A; B) corresponds to all events excluding the summer 2018 heatwave. The second row (C; D) instead, analysed the entire dataset. The first two principal components account for approximately 70-80% of the variance. Ellipses represent different sampling cruises. Other abbreviations: SSC, side scatter; FSC, forward scatter; FL2, FL3, FL4, photopigments composition and quantity (respectively detect: phycoerythrin, chlorophyll a and b, phycocyanin, see Table 1 for the list of parameters and abbreviations used). Arrows indicate the covariance between cruises and main physiological features describing community composition. The color gradient reports the sampling temperature at sea in the respective cruises (blue to red from colder to warmer temperatures).

**Figure. 5.**
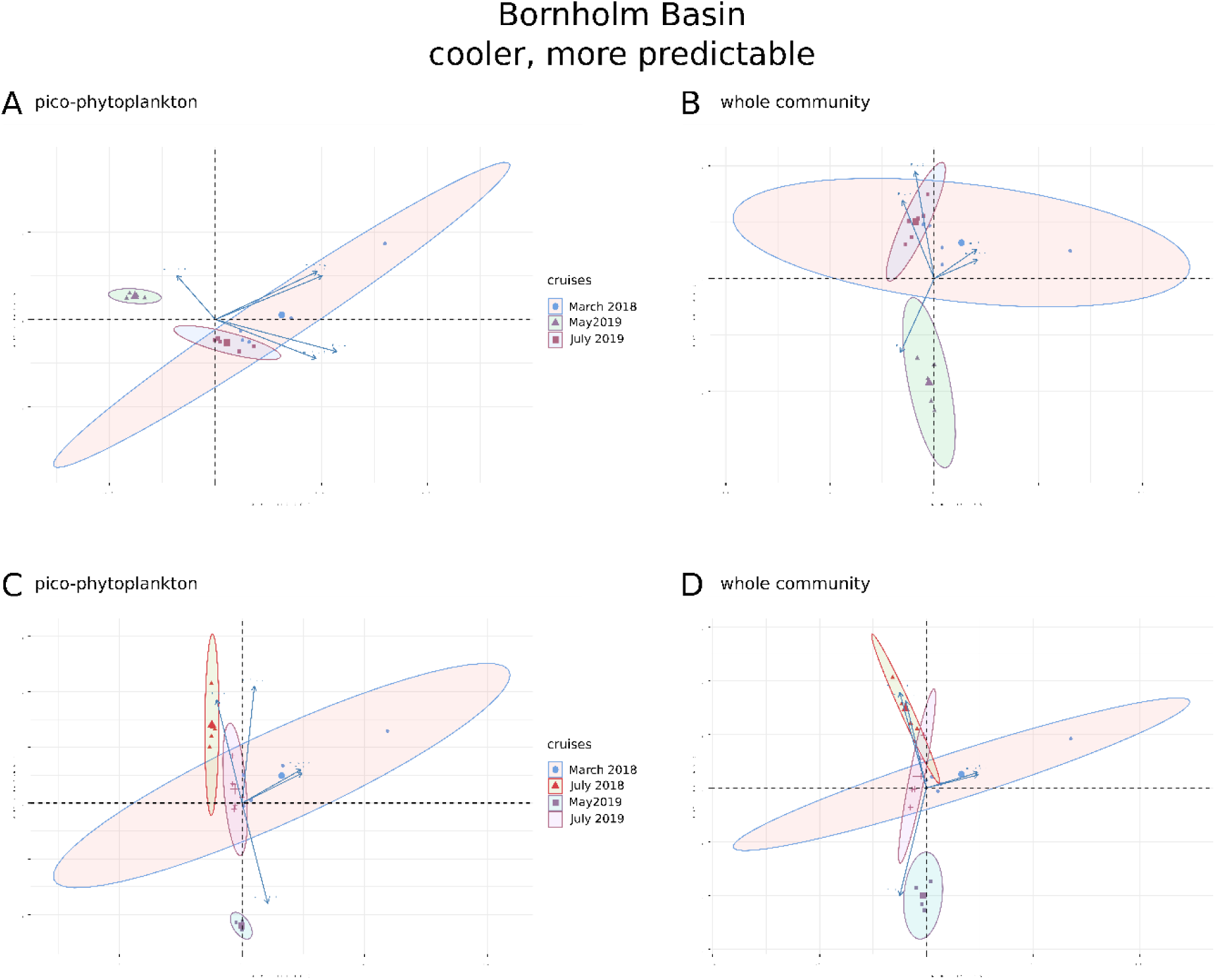
PCA of cytometric fingerprint covariance in the Bornholm Basin whole community and pico-phytoplankton fraction. Loadings represent phenotypic characteristics of functional groups derived from flow cytometry. The first row (A; B) corresponds to all events excluding the summer 2018 heatwave. The second row (C; D) instead, analysed the entire dataset. The first two principal components account for approximately 70-80% of the variance. Ellipses represent different sampling cruises. Other abbreviations: SSC, side scatter; FSC, forward scatter; FL2, FL3, FL4, photopigments composition and quantity (respectively detect: phycoerythrin, chlorophyll a and b, phycocyanin, see Table 1 for the list of parameters and abbreviations used). Arrows indicate the covariance between cruises and main physiological features describing community composition. The color gradient reports the sampling temperature at sea in the respective cruises (blue to red from colder to warmer temperatures).

Seasonal differences in the communities’ flow cytometric data were largely driven by changes in size and internal granularity of the cells (FSC and SSC respectively; Fig 5 B C D). Only when excluding the heatwave event, FL4 (phycocyanin content, a prominent photopigment in cyanobacteria) influenced the maximum variance direction in the data in some cases (i.e. Kiel Area whole community and pico-phytoplankton; Bornholm Basin pico-phytoplankton).

## Discussion

We investigated the direct effect of temperature on phytoplankton metabolic rates on physiological time scales (response curves) as well as ecological and evolutionary processes on longer timescales (seasonal differences, differences between basins and interactions thereof). We used a local approach of comparing adjacent but characteristically different sea surface areas and thereby minimised the confounding effects that arise when comparing regions across global scales.

We show that overall T_opt_GP in phytoplankton communities <35 µm strongly increases with sea surface temperatures, in contrast to T_opt_RESP, which is less sensitive to warming. Regions differed strongly in their sensitivity to heatwaves and in how community composition changed throughout the seasons. While in line with rapid thermal evolution in phytoplankton, this indicates differences in the underlying adaptive patterns due to variation with overarching fundamental and realised niches (i.e. the physical-chemical conditions and the biological interactions respectively (Hutchinson, 1957)). Similarly, in the absence of heatwaves, seasonality was the main driver for changes in T_opt_GP, indicating that thermal performances co-vary systematically with environmental parameters (Padfield *et al*, 2017). T_opt_ was overall the most reactive parameter in our dataset compared to the other parameters of the curve (i.e. E_a_, E_h_, lnc). This is in strong contrast with results from single species acclimated to changes in mean temperature under laboratory conditions, or model predictions. There, the shape of the curve changes rapidly, and parameters that change most with warming tend to be the elevation and steepness (E_a_ and E_h_, respectively) of the thermal response curve (Barton *et al*., 2018), and, on short time scales, respiration is more sensitive than photosynthesis.

As expected (Thomas *et al*., 2017) we found that in all samples from the more predictable, cooler Bornholm Basin, and in community samples <35µm from the warmer, more unpredictable Kiel Area, T_opt_GP was higher than the mean environmental temperature. More unexpectedly, in the pico-phytoplankton samples from the Kiel Area, sampling temperatures regularly exceeded T_opt_GP. There, T_opt_GP did not change with seasonal temperature, but remained stable at ca 16 °C throughout the year (for comparison, the five-year average temperature in the Kiel Area is 7.92 ºC), as depicted by 1:1 regression line in Fig. 2 and 3 and pairwise comparisons. Growth rates of small phytoplankton is often positively correlated with temperature (Kulk *et al*., 2012), so it would stand to reason that the photosynthetic traits that commonly underpin fitness (Cullen, 1990) behave in a similar fashion. Differences in responsiveness of T_opt_GP between the two size classes may be due to a buffering effect caused by a higher biodiversity, which is intuitively higher in the whole community fraction due to the contribution of cells larger than 3 µm, and in line with theoretical and empirical frameworks (e.g García *et al*., 2018). Moreover, cyanobacteria tolerate high temperature well, but they are more common in the Bornholm Basin rather than in the Kiel Area (Öberg, 2017).

Phytoplankton growth is a balance between GP and RESP, since a large part of the carbon produced by photosynthesis is then remineralised by respiration (Falkowski *et al*., 1998). Surprisingly, T_opt_ for respiration did not follow the seasonally increasing sea surface temperature at all. The lack of responsiveness of T_opt_RESP is unexpected (Staehr & Birkeland, 2006) and suggests phytoplankton communities could actively counterbalance seasonal increasing temperatures through adjusting photosynthetic activity alone. As on-board incubations to test whether this was reflected in growth rates across the same temperatures were not feasible, we cannot postulate direct effects on fitness based on this data set alone. While measures of Carbon Use Efficiency (CUE) could serve as an indirect measure of the amount of carbon available for growth (Gifford, 2003), we cannot with certainty estimate the contributions of heterotrophic bacterial respiration here, making calculations for CUE or NPP not completely reliable.

Under the influence of heatwaves (i.e. August 2018 cruise), the relationship between T_opt_RESP and sea surface temperatures greatly changed, creating a relationship that more closely aligns with model predictions concerning the slopes of thermal reaction norms, i.e. a higher reactivity of respiration than photosynthesis. When respiration rises with increasing temperatures, phytoplankton growth can be limited if respiration rises to a point where it exceeds photosynthesis. Hence, heatwaves have the potential to drastically alter phytoplankton primary production, especially in regions that are cooler and less variable, such as the Bornholm Basin.

Most recent studies on heat wave scenarios focused on their effect on community composition (e.g. Striebel *et al*., 2016). Several observations indicate that biodiversity is consistently lower during heatwaves (especially true for nutrient-poor regions (Hayashida *et al*., 2020) resulting in drastically changed community structures. Based on flow cytometric data, we found no evidence of the heatwave significantly reducing phenotypic diversity although the capacity of the communities to metabolically compensate for extreme and sudden events seemed to decline.

Crucially, our study shows that changes in thermal reaction norms can be rapid (on a seasonal time scale) enough to adjust to seasonal variability even when the selection environment is highly complex, but not to sudden events like heatwaves. Models assume that the degree to which plasticity evolves in a fluctuating environment is ultimately limited by the cost of plasticity, though the cost has remained elusive in experiments (Murren *et al*., 2015). If plasticity has an intrinsic cost, heterogeneous conditions might avoid a perfect match between phenotype and environment (i.e. mean phenotype is the optimum phenotype in all scenarios) and instead force a generalist/specialist coupling. In the latter case, organisms will perform quite well in a variety of environments, but not quite as well as in the specialist specific niche (Angilletta *et al*., 2003). We found that regardless of the variability and fluctuations’ complexity of the previously experienced environment, i.e. the geographical region, all the analysed samples reacted the same way and we detected neither specialistic nor generalistic trends. Even if we had conducted our investigation under more natural conditions (i.e. minimising acclimation bias and forced biotic relationships), we still investigated only one environmentally induced pair of traits. Ideally, more traits (e.g. growth rates, oxidative stress) should have been examined since plasticity likely affected different traits with in theory different costs (e.g. Walworth *et al*., 2021). In our case, in the Kiel Area, populations’ reaction norms evolved closer to the optimum even in extreme conditions like during a heatwave. We can argue that the more unpredictable Kiel Area represents a not canalised environment, without decreased genetic and phenotypic variance. Therefore, maintenance costs, needed to track environmental conditions and changes, were here minimized (DeWitt, 1998).

On seasonal timescales, various mechanisms, such as phenotypic plasticity or biotic filtering (i.e. species less adapted to the conditions they face are outcompeted by more adapted ones (Thomas *et al*., 2016), can act on and determine the way communities respond to environmental changes. In our framework, changes on the functional group level are not an underlying reason for changes in thermal tolerance in the more variable Kiel Area. It is likely that communities adjust through plasticity or rapid evolutionary responses (including sorting of standing genetic variation of different genotypes within the same species). Populations in the Kiel Area were rather similar throughout cruises demonstrating that peculiar sensitivities of communities rather than functional traits, lead to adjustments of thermal tolerances. In the more predictable Bornholm Basin, the opposite situation occurred, with a strong shift on the functional group level. Ecological implications are grand and intuitive: our findings point out that, moving toward a generally warmer and more unpredictable future, changes of the communities’ composition in formerly predictable areas, might disrupt ecosystem functions. Populations in the Kiel Area had centuries to evolve to variable conditions, whereas current changes are happening on a much faster timescale.

Here we show that even in a complex natural environment, the ways in which major metabolic pathways react to changes in temperature are at least partially predictable (i.e. as temperature rises, so does photosynthetic T_opt_) and repeatable (our findings hold across size classes and regions). Although we currently lack information on the mechanistic processes involved in the maintenance of these patterns on acclimation timescales, our results provide essential information (e.g. timing of adaptive patterns, differential responses of natural assemblages) on adaptive dynamics of phytoplankton for ecosystem models. A better understanding of the timescales at which fast reproducing organisms react to changes could indeed make the coupling between human-induced changes and natural adaptive patterns more precisely predictable.

## References

Andrade-Restrepo, M., Champagnat, N. and Ferrière, R. (2019) ‘Local adaptation, dispersal evolution, and the spatial eco-evolutionary dynamics of invasion’, Ecology Letters, 22(5), pp. 767–777. doi: 10.1111/ele.13234.

Barton, S. et al. (2018) ‘Universal metabolic constraints on the thermal tolerance of marine phytoplankton’, bioRxiv. doi: 10.1101/358002.

Barton, S. et al. (2020) ‘Evolutionary temperature compensation of carbon fixation in marine phytoplankton’, Ecology Letters, 23(4), pp. 722–733. doi: 10.1111/ele.13469.

Bennett, A. F. and Lenski, R. E. (2007) ‘An experimental test of evolutionary trade-offs during temperature adaptation’, Proceedings of the National Academy of Sciences, 104(suppl 1), pp. 8649 LP–8654. doi: 10.1073/pnas.0702117104.

Bestion, E. et al. (2018) ‘Metabolic traits predict the effects of warming on phytoplankton competition’, Ecology Letters. Blackwell Publishing Ltd, pp. 655–664. doi: 10.1111/ele.12932.

Bopp, L. et al. (2013) ‘Multiple stressors of ocean ecosystems in the 21st century: projections with CMIP5 models’, Biogeosciences, 10(10), pp. 6225–6245. doi: 10.5194/bg-10-6225-2013.

Botero, C. A. et al. (2015) ‘Evolutionary tipping points in the capacity to adapt to environmental change’, Proceedings of the National Academy of Sciences of the United States of America, 112(1), pp. 184–189. doi: 10.1073/pnas.1408589111.

Boyce, D. G., Lewis, M. R. and Worm, B. (2010) ‘Global phytoplankton decline over the past century’, Nature, 466(7306), pp. 591–596. doi: 10.1038/nature09268.

Boyd, P. W. et al. (2016) ‘Biological responses to environmental heterogeneity under future ocean conditions’, Global change biology, 22(8), pp. 2633–2650. doi: 10.1111/gcb.13287.

Brown, J. H. et al. (2004) ‘Perspectives’, Ecology, 85(7), pp. 1771–1789.

Carr, M. H. et al. (2016) ‘Comparing Marine and Terrestrial Ecosystems : Implications for the Design of Coastal Marine Reserves Warner and John L. Largier Source : Ecological Applications, Vol. 13, No. 1, Supplement : The Science of Marine Reserves Published by : Wiley Stable’, 13(1).

Cullen, J. J. (1990) ‘On models of growth and photosynthesis in phytoplankton’, Deep Sea Research Part A. Oceanographic Research Papers, 37(4), pp. 667–683. doi: https://doi.org/10.1016/0198-0149(90)90097-F.

Dell, A. I., Pawar, S. and Savage, V. M. (2011) ‘Systematic variation in the temperature dependence of physiological and ecological traits’, Proceedings of the National Academy of Sciences, 108(26), pp. 10591 LP–10596. doi: 10.1073/pnas.1015178108.

DeWitt, T. J. (1998) ‘Costs and limits of phenotypic plasticity: Tests with predator-induced morphology and life history in a freshwater snail’, Journal of Evolutionary Biology. John Wiley & Sons, Ltd, 11(4), pp. 465–480. doi: https://doi.org/10.1046/j.1420-9101.1998.11040465.x.

Eilers, P. H. C. and Peeters, J. C. H. (1988) A MODEL FOR THE RELATIONSHIP BETWEEN LIGHT. INTENSITY AND THE RATE OF PHOTOSYNTHESIS IN PHYTOPLANKTON, Ecological Modelling.

Eppley, R. W. (1972) ‘Fishery Bullettin, Volume 70, Issue 4’, in Fishery Bulletin, p. 1063.

Falkowski, P. G., Barber, R. T. and Smetacek, V. (1998) ‘Biogeochemical Controls and Feedbacks on Ocean Primary Production’, Science, 281(5374), pp. 200 LP–206. doi: 10.1126/science.281.5374.200.

Field, C. B. et al. (1998) ‘Primary Production of the Biosphere: Integrating Terrestrial and Oceanic Components’, Science, 281(5374), pp. 237 LP–240. doi: 10.1126/science.281.5374.237.

García, F. C. et al. (2018) ‘Changes in temperature alter the relationship between biodiversity and ecosystem functioning’, Proceedings of the National Academy of Sciences of the United States of America, 115(43), pp. 10989–10994. doi: 10.1073/pnas.1805518115.

Gifford, R. M. (2003) ‘Plant respiration in productivity models: conceptualisation, representation and issues for global terrestrial carbon-cycle research’, Functional Plant Biology, 30(2), pp. 171–186. Available at: https://doi.org/10.1071/FP02083.

del Giorgio, P. A., Cole, J. J. and Cimbleris, A. (1997) ‘Respiration rates in bacteria exceed phytoplankton production in unproductive aquatic systems’, Nature, 385(6612), pp. 148–151. doi: 10.1038/385148a0.

Godhe, A. and Rynearson, T. (2017) ‘The role of intraspecific variation in the ecological and evolutionary success of diatoms in changing environments’, Philosophical Transactions of the Royal Society B: Biological Sciences, 372(1728). doi: 10.1098/rstb.2016.0399.

Grasshoff, K., Kremling, K. and Ehrhardt, M. (2009) Methods of Seawater Analysis. Wiley-VCH.

Hayashida, H., Matear, R. J. and Strutton, P. G. (2020) ‘Background nutrient concentration determines phytoplankton bloom response to marine heatwaves’, Global Change Biology. John Wiley & Sons, Ltd, 26(9), pp. 4800–4811. doi: https://doi.org/10.1111/gcb.15255.

Hobday, A. J. et al. (2016) ‘A hierarchical approach to defining marine heatwaves’, Progress in Oceanography. Elsevier Ltd, 141, pp. 227–238. doi: 10.1016/j.pocean.2015.12.014.

Hoppe, C. J. M. et al. (2018) ‘Compensation of ocean acidification effects in Arctic phytoplankton assemblages’, Nature Climate Change. Nature Publishing Group, 8(6), pp. 529–533. doi: 10.1038/s41558-018-0142-9.

Hutchinson, G. E. (1957) ‘Population studies-animal ecology and demography-concluding remarks’, in Cold Spring Harbor symposia on quantitative biology. Cold Spring Harbor, NY: Cold Spring Harbor lab press, Pubblications dept.

Jr, M. J. A. et al. (2003) ‘Tradeoffs and the evolution of thermal reaction norms’, 18(5), pp. 234–240. doi: 10.1016/S0169-5347(03)00087-9.

Karl, D. M. et al./person-group>. (2003) ‘Temporal Studies of Biogeochemical Processes Determined from Ocean Time-Series Observations During the JGOFS Era’, in Fasham, M. J. R. (ed.) Ocean Biogeochemistry: The Role of the Ocean Carbon Cycle in Global Change. Berlin, Heidelberg: Springer Berlin Heidelberg, pp. 239–267. doi: 10.1007/978-3-642-55844-3_11.

Karl, T. R. and Trenberth, K. E. (2003) ‘Modern Global Climate Change’, Science, 302(5651), pp. 1719–1723. doi: 10.1126/science.1090228.

Kingsolver, J. G. (2009) ‘The Well-Temperatured Biologist (American Society of Naturalists Presidential Address) *’, 174(6). doi: 10.1086/648310.

Kirkwood, D. (1996) ‘Nutrients: practical notes on their determination in seawater’, ICES Techniques in Marine Environmental Sciences, (17), p. 23 pp.

Kulk, G. et al. (2012) ‘Temperature-dependent growth and photophysiology of prokaryotic and eukaryotic oceanic picophytoplankton’, Marine Ecology Progress Series, 466, pp. 43–55. doi: 10.3354/meps09898.

Lande, R. (2014) ‘Evolution of phenotypic plasticity and environmental tolerance of a labile quantitative character in a fluctuating environment’, Journal of Evolutionary Biology, 27(5), pp. 866–875. doi: 10.1111/jeb.12360.

Laufkotter, C. et al. (2015) ‘Drivers and uncertainties of future global marine primary production in marine ecosystem models’, Biogeosciences, 12(23), pp. 6955–6984. doi: 10.5194/bg-12-6955-2015.

Lee, C. K. and Peterson, Æ. M. E. (2008) ‘The effect of temperature on enzyme activity : New insights and their implications The effect of temperature on enzyme activity : new insights and their implications’, (February 2014). doi: 10.1007/s00792-007-0089-7.

López-Urrutia, Á. et al. (2006) ‘Scaling the metabolic balance of the oceans’, Proceedings of the National Academy of Sciences, 103(23), pp. 8739 LP–8744. doi: 10.1073/pnas.0601137103.

Magozzi, S. and Calosi, P. (2015) ‘Integrating metabolic performance, thermal tolerance, and plasticity enables for more accurate predictions on species vulnerability to acute and chronic effects of global warming’, Global Change Biology, 21(1), pp. 181–194. doi: 10.1111/gcb.12695.

Mitchell, S. E. and Lampert, W. (2000) ‘Temperature adaptation in a geographically widespread zooplankter, Daphnia magna’, 13, pp. 371–382.

Morán, X. A. G. et al. (2010) ‘Increasing importance of small phytoplankton in a warmer ocean’, Global Change Biology, 16(3), pp. 1137–1144. doi: 10.1111/j.1365-2486.2009.01960.x.

Munguia, P. and Alenius, B. (2013) ‘The role of preconditioning in ocean acidification experiments: a test with the intertidal isopod Paradella dianae’, Marine and Freshwater Behaviour and Physiology. Taylor & Francis, 46(1), pp. 33–44. doi: 10.1080/10236244.2013.788287.

Murphy, J. and Riley, J. P. (1962) ‘A Modified Single Solution Method for the Determination of Phosphate in Natural Waters’, Analytica Chimica Acta, 27, pp. 31–36. doi: 10.1057/9781137461131.

Murren, C. J. et al. (2015) ‘Constraints on the evolution of phenotypic plasticity: limits and costs of phenotype and plasticity’, Heredity, 115(4), pp. 293–301. doi: 10.1038/hdy.2015.8.

Öberg, J. (2017) ‘Cyanobacteria blooms in the Baltic Sea’. HELCOM Baltic Sea Environment Fact Sheets.

Pacifici, M. et al. (2015) ‘Assessing species vulnerability to climate change’, Nature Climate Change, 5(3), pp. 215–224. doi: 10.1038/nclimate2448.

Padfield, D. et al. (2016) ‘Rapid evolution of metabolic traits explains thermal adaptation in phytoplankton’, Ecology Letters, 19(2), pp. 133–142. doi: 10.1111/ele.12545.

Padfield, D. et al. (2017) ‘Metabolic compensation constrains the temperature dependence of gross primary production’, Ecology Letters, 20(10), pp. 1250–1260. doi: 10.1111/ele.12820.

Pawar, S., Dell, A. I. and Savage, V. M. (2015) ‘Chapter 1 - From Metabolic Constraints on Individuals to the Dynamics of Ecosystems’, in Belgrano, A., Woodward, G., and Jacob, U. B. T.-A. F. B. (eds). San Diego: Academic Press, pp. 3–36. doi: https://doi.org/10.1016/B978-0-12-417015-5.00001-3.

Pörtner, H. O. (2002) ‘Climate variations and the physiological basis of temperature dependent biogeography: systemic to molecular hierarchy of thermal tolerance in animals’, Comparative Biochemistry and Physiology Part A: Molecular & Integrative Physiology, 132(4), pp. 739–761. doi: https://doi.org/10.1016/S1095-6433(02)00045-4.

Post, D. M. and Palkovacs, E. P. (2009) ‘Eco-evolutionary feedbacks in community and ecosystem ecology: interactions between the ecological theatre and the evolutionary play’, Philosophical Transactions of the Royal Society B: Biological Sciences. Royal Society, 364(1523), pp. 1629–1640. doi: 10.1098/rstb.2009.0012.

Regaudie-de-Gioux, A. et al. (2014) ‘Comparing marine primary production estimates through different methods and development of conversion equations‘, Frontiers in Marine Science, p. 19. Available at: https://www.frontiersin.org/article/10.3389/fmars.2014.00019.

Schaum, C. E. et al. (2017) ‘Adaptation of phytoplankton to a decade of experimental warming linked to increased photosynthesis’, Nature Ecology and Evolution. Nature Publishing Group, 1(4). doi: 10.1038/s41559-017-0094.

Schoolfield, R. M., Sharpe, P. J. H. and Magnuson, C. E. (1980) ‘Non-linear Regression of Biological Temperature-dependent Rate Models Based on Absolute Reaction-rate Theory’.

Soeijs-Leijonmalm, P. and Andrén, E. (2017) ‘No Title’, in Biological Oceanography of the Baltic Sea. Springer Science+Business Media Dordrecht, pp. 23–84.

Souther, S. and Graw, J. B. M. C. (2011) ‘Evidence of Local Adaptation in the Demographic Response of American Ginseng to Interannual Temperature Variation Abstract ‘:, 25(5), pp. 922–931. doi: 10.1111/j.1523-1739.2011.01695.x.

Staehr, P. A. and Birkeland, M. J. (2006) ‘Temperature acclimation of growth, photosynthesis and respiration in two mesophilic phytoplankton species’, Phycologia, 45(6), pp. 648–656. doi: 10.2216/06-04.1.

Starrfelt, J. and Kokko, H. (2012) ‘Bet-hedging—a triple trade-off between means, variances and correlations’, Biological Reviews. John Wiley & Sons, Ltd, 87(3), pp. 742–755. doi: 10.1111/j.1469-185X.2012.00225.x.

Striebel, M. et al. (2016) ‘Phytoplankton responses to temperature increases are constrained by abiotic conditions and community composition’, Oecologia, 182(3), pp. 815–827. doi: 10.1007/s00442-016-3693-3.

Thomas, M. et al. (2012) ‘A global pattern of thermal adatpation in marine phytoplankton.’, Science, 338(338), pp. 1085–1088. Available at: http://science.sciencemag.org/content/sci/338/6110/1085.full.pdf.

Thomas, M. K. et al. (2017) ‘Temperature–nutrient interactions exacerbate sensitivity to warming in phytoplankton’, Global Change Biology, 23(8), pp. 3269–3280. doi: 10.1111/gcb.13641.

Thomas, M. K., Kremer, C. T. and Litchman, E. (2016) ‘Environment and evolutionary history determine the global biogeography of phytoplankton temperature traits’, Global Ecology and Biogeography. Blackwell Publishing Ltd, 25(1), pp. 75–86. doi: 10.1111/geb.12387.

Visser, P. M. et al. (2016) ‘How rising CO2 and global warming may stimulate harmful cyanobacterial blooms’, Harmful Algae, 54, pp. 145–159. doi: https://doi.org/10.1016/j.hal.2015.12.006.

Walworth, N. G. et al. (2021) ‘The evolution of trait correlations constrains phenotypic adaptation to high CO2 in a eukaryotic alga’, Proceedings of the Royal Society B: Biological Sciences. Royal Society, 288(1953), p. 20210940. doi: 10.1098/rspb.2021.0940.

Wolf, K. K. E., Hoppe, C. J. M. and Rost, B. (2018) ‘Resilience by diversity: Large intraspecific differences in climate change responses of an Arctic diatom’, Limnology and Oceanography. Wiley Blackwell, 63(1), pp. 397–411. doi: 10.1002/lno.10639.

Worden, A. Z. and Not, F. (2008) ‘Ecology and diversity of picoeukaryotes’, in Kirchman, D. L. (ed.) Microbial Ecology of the Oceans. Second Edi. John Wiley & Sons, Inc., pp. 159–205.

Zhong, D. et al. (2020) ‘Functional redundancy in natural pico-phytoplankton communities depends on temperature and biogeography’, bioRxiv, p. 2020.05.14.096123. doi: 10.1101/2020.05.14.096123

