## Supplementary information for "Predicting the unpredictable: heatwaves and history of variability shape phytoplankton community thermal responses within one generation"


| | | |
|---|---|---|
| **Table S1** | Analysis of abiotic factors at sampling site | Page 2 |
| **Table S2** | Ranges of temperatures used for metabolism´s responses | Page 2 |
| **Table S3** | Analysis of decomposition analysis´ random component | Page 3 |
| **Table S4** | Model selection for gross photosynthesis and respiration $T_{opt}$ | Page 8 |
| **Table S5** | PerMANOVA results on cytometric fingerprints | Page 10 |
| **Figure S1** | Decomposition analysis of thermal profiles of the two geographical areas (Kiel Area and Bornholm Basin) | Page 3 |
| **Figure S2** | Nutrients´ concentrations in the two geographical areas | Page 6 |
| **Figure S3** | Relative abundances of pico-phytoplankton and whole community | Page 7 |

**Table S1.**Statistical results from a One-way Anova using the “Anova” function in the *car* package on R. The test was conducted on means for the main abiotic factors (salinity, temperature and nutrient concentration) describing the Kiel Basin and the Mecklenburger Bight. No significant differences were present, and we therefore group the two regions as ‘Kiel Area’ throughout the paper. (df: degree of freedom; SS: sum of squares; F: F-value; p: p-value).

| Variable | *df* | SS | *F* | *p* |
|---|---|---|---|---|
| Salinity (PSU) | 1 | 22.038 | 3.521 | 0.085 |
| Temperature (°C) | 1 | 0.04 | 6e-04 | 0.98 |
| Nitrogen | 1 | 312.2 | 0.251 | 0.627 |
| Silicates | 1 | 1.421 | 0.11 | 0.747 |
| Phosphate | 1 | 47.84 | 0.483 | 0.503 |

**Table S2.** Ranges of temperatures used for metabolism analysis. Increments are uneven and higher temperature may vary due to tolerance differences of the samples.

| | Temperature (°C) | | | | | | | | | | | | | | | | | | | | | |
|---|---|---|---|---|---|---|---|---|---|---|---|---|---|---|---|---|---|---|---|---|---|---|
| **Cruise id** | **3** | **5** | **8** | **10** | **12** | **14** | **15** | **18** | **20** | **22** | **24** | **25** | **26** | **28** | **29** | **30** | **31** | **32** | **34** | **35** | **38** | **40** |
| AL505 (March 2018) | ✓ | ✓ | | ✓ | ✓ | | ✓ | ✓ | ✓ | | | ✓ | | | | | | | | | | |
| AL513 (July 2018) | | | | ✓ | | ✓ | | ✓ | | ✓ | | | ✓ | | | ✓ | | | ✓ | | ✓ | ✓ |
| AL520 (March 2019) | ✓ | ✓ | ✓ | ✓ | ✓ | | ✓ | ✓ | ✓ | ✓ | | ✓ | | | | | | | | | | |
| AL521 (April 2019) | | ✓ | ✓ | ✓ | ✓ | | ✓ | ✓ | ✓ | ✓ | | | | | | | | | | | | |
| AL522 (May 2019) | | ✓ | ✓ | ✓ | ✓ | | ✓ | ✓ | ✓ | ✓ | ✓ | | ✓ | | | | | | | | | |
| AL524 (July 2019) | | | ✓ | ✓ | ✓ | | ✓ | ✓ | ✓ | | | ✓ | | | ✓ | | ✓ | ✓ | | ✓ | | |
| AL530 (October 2019) | | ✓ | ✓ | ✓ | ✓ | | ✓ | ✓ | ✓ | ✓ | ✓ | | ✓ | ✓ | | | | | | | | |

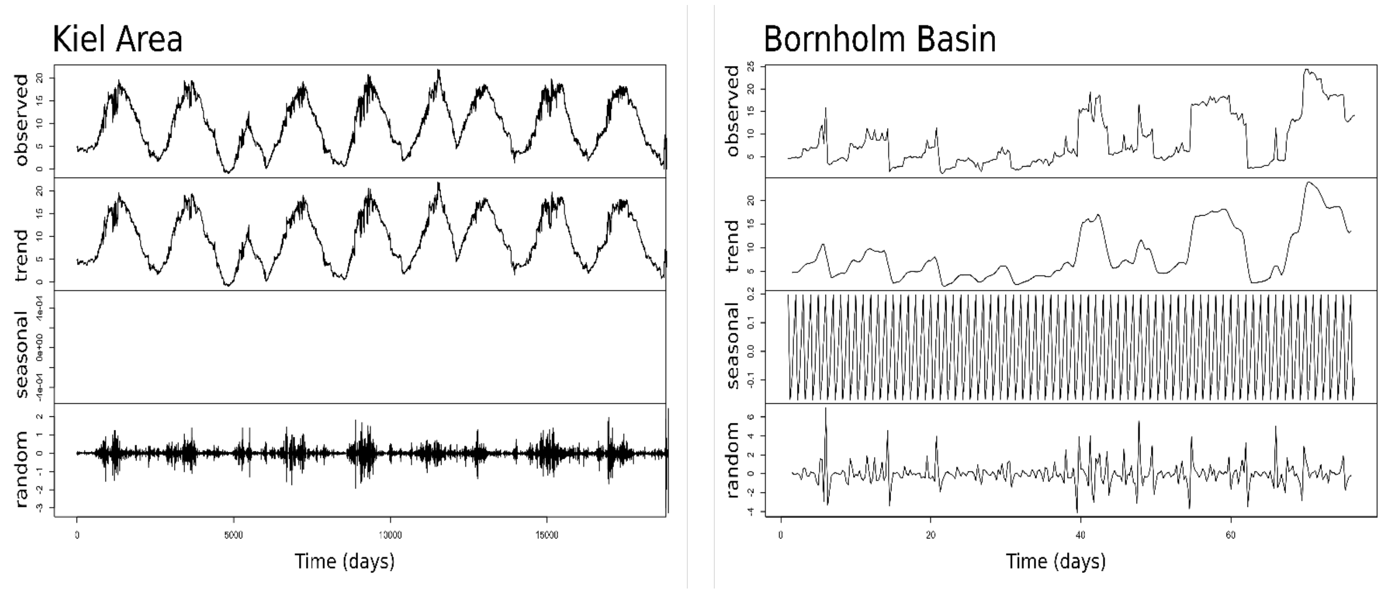


**Figure S1.** Decomposition analyses of surface temperatures (-8 m) of the last 5 years for the Kiel Area and the Bornholm Basin. Absence of a reproducible and clear pattern of seasonality in the Kiel Area, indicates a less predictable trend in the area.

**Table S3.** Random components outcomes produced from decomposition analysis performed on sea surface temperatures time series for Bornholm Basin and the Kiel Area, using the function decompose of the anomalize package (0.2.0). We used an additive (seasonal + trend + random) approach, assuming a quarterly seasonality (frequency = 4. The quarterlies are reported in the table as Qtr1, Qtr2, Qtr 3 and Qtr4). Statistical results from a One-way ANOVA comparing the two geographical areas are reported at the beginning (df: degree of freedom; SS: sum of squares; F: F-value; p: p-value). Mean values for random effect were higher in the Kiel Area, meaning that the time series is less constant and consequently more variable (KA: 1202.95 ± 682.67; BB: 151.94 ± 87.47). No seasonal component was found for the Kiel Area (also using a multiplicative approach assuming a monthly seasonality).

| Variable | *df* | SS | *F* | *p* |
|---|---|---|---|---|
| Geographical Area | 1 | 3.001e+08 | 718.9 | >2e-16 |

**Bornholm Basin**

| Time points | Qtr1 | Qtr2 | Qtr3 | Qtr4 |
|---|---|---|---|---|
| 1 | NA | NA | 0.122195559 | 0.239398849 |
| 2 | 0.110243914 | -0.464088322 | -0.154304441 | -0.316601151 |
| 3 | 0.707993914 | 0.055411678 | -0.260179441 | 0.055711349 |
| 4 | 0.352743914 | -0.387150822 | 0.023695559 | -0.455913651 |
| 5 | -1.074443586 | 1.117911678 | 2.453695559 | -0.941913651 |
| 6 | 1.572181414 | -1.914775822 | -0.814804441 | 0.118898849 |
| 7 | 0.171368914 | -0.377838322 | 0.014320559 | 0.310711349 |
| 8 | 0.104118914 | -0.117088322 | 0.101258059 | -0.628351151 |
| 9 | -0.999631086 | 1.091974178 | 0.735508059 | -0.041726151 |
| 10 | 0.086743914 | -0.714400822 | 0.297820559 | -0.145788651 |

| | | | | |
|---|---|---|---|---|
| 11 | -0.502506086 | -0.827588322 | 1.854320559 | 0.633086349 |
| 12 | -0.887943586 | -0.967025822 | 1.688508059 | -0.330101151 |
| 13 | -0.547693586 | -0.623400822 | 1.216445559 | -0.360976151 |
| 14 | -0.049881086 | 4.050161678 | -3.393616941 | -0.806601151 |
| 15 | 0.262868914 | -0.470963322 | 0.140883059 | 0.274273849 |
| 16 | -0.268068586 | -0.948900822 | 0.766633059 | 0.371273849 |
| 17 | -0.001193586 | -0.104025822 | -0.128929441 | 0.367273849 |
| 18 | 0.052868914 | -0.460213322 | 0.053508059 | -0.120601151 |
| 19 | -0.160631086 | -0.914838322 | 1.822445559 | -0.089851151 |
| 20 | -0.342943586 | -0.446963322 | -0.224616941 | 2.462898849 |
| 21 | 0.952675164 | -2.115238322 | -0.895104441 | 0.024586349 |
| 22 | 0.282212664 | -0.296225822 | -0.003866941 | 0.265398849 |
| 23 | -0.062381086 | -0.987088322 | 0.690008059 | 0.608648849 |
| 24 | -0.094881086 | -0.329838322 | -0.051304441 | 0.133773849 |
| 25 | 0.391493914 | -0.289775822 | 0.208633059 | 0.326336349 |
| 26 | -0.579506086 | 0.427286678 | -0.334429441 | -0.791476151 |
| 27 | 0.590993914 | -0.146963322 | 0.065445559 | 0.191336349 |
| 28 | 0.044118914 | -0.287025822 | -0.068929441 | -0.339476151 |
| 29 | 0.581368914 | -0.543713322 | 1.052695559 | -0.069851151 |
| 30 | -0.604881086 | 0.636224178 | 1.335570559 | -1.088913651 |
| 31 | -0.357756086 | -0.441900822 | 0.044758059 | 0.135836349 |
| 32 | -0.148506086 | -0.112838322 | 0.064445559 | 0.327148849 |
| 33 | -0.116443586 | -0.709213322 | 0.416820559 | 0.473148849 |
| 34 | -0.310381086 | -0.176588322 | 0.356695559 | 0.309836349 |
| 35 | -0.598443586 | 0.143786678 | 0.512070559 | -0.493726151 |
| 36 | -0.274631086 | -0.839838322 | 1.243570559 | 0.217398849 |
| 37 | -0.161756086 | -0.483838322 | -0.447491941 | -0.646288651 |
| 38 | 2.714681414 | -0.878900822 | -0.463179441 | -0.089788651 |
| 39 | 0.782556414 | -1.649025822 | -4.133429441 | 4.272836349 |
| 40 | 0.690868914 | 0.046536678 | -0.003679441 | 0.096648849 |
| 41 | -1.839506086 | 1.859036678 | 2.977570559 | -2.390851151 |
| 42 | -2.989506086 | 1.615286678 | 0.696320559 | 0.377898849 |
| 43 | 2.079243914 | -0.822213322 | -0.791179441 | 0.164461349 |
| 44 | 2.250743914 | -2.525463322 | -0.659804441 | -0.013163651 |
| 45 | 0.212681414 | -0.408338322 | 0.015195559 | -0.108351151 |
| 46 | -0.962318586 | 2.329536678 | -0.927179441 | 0.082961349 |
| 47 | -0.283568586 | 0.558786678 | -0.664929441 | -0.890913651 |
| 48 | -3.284318586 | 5.379411678 | 0.797070559 | -1.172101151 |
| 49 | -0.389506086 | -0.634713322 | 0.027570559 | 0.676148849 |
| 50 | 2.856618914 | -2.427025822 | -0.767429441 | 0.167836349 |
| 51 | -0.033881086 | 0.023099178 | -0.197179441 | 0.019711349 |
| 52 | -0.048256086 | -0.097213322 | -0.059929441 | -0.059601151 |
| 53 | -0.131193586 | 0.658224178 | -0.813429441 | 0.275398849 |
| 54 | 1.223118914 | -0.848650822 | -0.005554441 | -1.229726151 |
| 55 | -3.858068586 | 3.690161678 | 1.161820559 | 0.415398849 |
| 56 | -0.014506086 | -0.634713322 | 0.158820559 | 0.640398849 |
| 57 | 0.022993914 | -1.034713322 | -0.053679441 | 1.052898849 |
| 58 | -0.252006086 | 0.202786678 | -0.541179441 | 0.040398849 |
| 59 | 0.329243914 | -0.428463322 | 0.071320559 | 0.465398849 |
| 60 | 0.047993914 | -0.197213322 | 2.283820559 | -1.997101151 |

**Kiel Area**

| Time points | Qtr1 | Qtr2 | Qtr3 | Qtr4 |
|---|---|---|---|---|
| 1 | NA | NA | 0.1501879117 | -0.0414275030 |
| 2 | -0.1850069240 | -0.0138159846 | -0.0983120883 | 0.0445724970 |
| 3 | 0.0692430760 | -0.0346284846 | -0.1024995883 | 0.0576974970 |
| 4 | 0.0686180760 | -0.0298784846 | -0.0514995883 | -0.0048025030 |
| 5 | -0.0518194240 | 0.0516215154 | 0.1470629117 | -0.0595525030 |
| 6 | -0.0146944240 | 0.0221215154 | -0.0716870883 | 0.0135724970 |
| 7 | -0.0296319240 | 0.1144965154 | -0.0788120883 | -0.0411775030 |
| 8 | 0.0273680760 | 0.0908715154 | 0.0528129117 | -0.0043025030 |
| 9 | -0.0236319240 | 0.0149965154 | -0.0714370883 | -0.1023025030 |
| 10 | -0.0007569240 | 0.0528715154 | 0.0075629117 | -0.0033025030 |
| 11 | 0.0016180760 | 0.0333715154 | 0.0158129117 | -0.0014275030 |
| 12 | 0.0433680760 | 0.0303715154 | 0.0039379117 | -0.1061775030 |
| 13 | -0.0153819240 | 0.0504965154 | -0.0601870883 | -0.1453025030 |
| 14 | 0.0199930760 | 0.0281215154 | 0.0334379117 | -0.0116775030 |
| 15 | 0.0176180760 | 0.1341215154 | -0.0663745883 | -0.0459900030 |
| 16 | 0.0499930760 | 0.0196215154 | 0.0032504117 | 0.0301349970 |
| 17 | -0.0341319240 | -0.0130034846 | -0.0115620883 | -0.0605525030 |
| 18 | -0.0178819240 | 0.0238715154 | 0.0470629117 | 0.0209474970 |
| 19 | 0.0043680760 | 0.0186215154 | -0.0199370883 | -0.0260525030 |
| 20 | 0.0212430760 | 0.0044965154 | 0.0216879117 | 0.0301974970 |
| 21 | 0.0022430760 | 0.0486215154 | 0.0004379117 | -0.0446775030 |
| 22 | -0.0653819240 | -0.0140034846 | -0.0253120883 | -0.0005525030 |
| 23 | -0.0516319240 | -0.0021284846 | 0.0456879117 | 0.0778849970 |
| 24 | -0.1276319240 | -0.0991284846 | 0.3184379117 | -0.1776150030 |
| 25 | -0.0646319240 | 0.1593715154 | 0.0635629117 | -0.1306775030 |
| 26 | -0.0940069240 | 0.0672465154 | -0.0588120883 | 0.1370099970 |
| 27 | -0.2221319240 | -0.0476284846 | 0.0931254117 | 0.1151349970 |
| 28 | 0.0109930760 | -0.1062534846 | -0.1401245883 | 0.1551974970 |
| 29 | 0.0139930760 | -0.0377534846 | -0.0246245883 | 0.0885724970 |
| 30 | 0.0649305760 | 0.0836840154 | -0.0731870883 | -0.1101775030 |
| 31 | -0.0060694240 | -0.0728159846 | 0.0291254117 | 0.2403224970 |
| 32 | -0.0641319240 | -0.3026909846 | -0.2269370883 | 0.3764474970 |
| 33 | -0.0027569240 | 0.0843715154 | 0.2526254117 | -0.4258025030 |
| 34 | 0.3583680760 | 0.6531215154 | -1.0286245883 | 0.0111349970 |
| 35 | -0.0467569240 | -0.2799409846 | -0.1325620883 | -0.0846775030 |
| 36 | 0.2113680760 | 0.8166215154 | 0.1023129117 | -0.0884900030 |
| 37 | 0.2693055760 | -0.1147534846 | -0.7823120883 | -0.1128025030 |
| 38 | 0.3785555760 | 0.0572465154 | -0.3061870883 | 0.2485099970 |
| 39 | -0.0381319240 | -0.1323159846 | -0.0834370883 | 0.3056349970 |
| 40 | -0.0375069240 | 0.0130590154 | -0.1800620883 | -0.1345525030 |
| 41 | -0.2428819240 | 0.7088715154 | 0.2513129117 | -0.1168025030 |
| 42 | -0.1827569240 | 0.0041215154 | -0.0965620883 | -0.1425525030 |
| 43 | 0.1004930760 | 0.1007465154 | -0.0121870883 | -0.0575525030 |
| 44 | 0.0057430760 | 0.1002465154 | 0.1120004117 | -0.0971775030 |
| 45 | 0.2220555760 | -0.5903784846 | 0.1380004117 | 0.3321974970 |
| 46 | 0.9124305760 | 0.0132465154 | -0.2468120883 | -0.4562400030 |
| 47 | -0.7778819240 | -0.5606284846 | 0.2115004117 | 0.6186974970 |
| 48 | -0.5977569240 | 0.3321215154 | 0.3550004117 | 0.8751974970 |
| 49 | 0.4993680760 | -1.4892534846 | -1.0240620883 | 0.3529474970 |
| 50 | 0.3849930760 | 0.2703090154 | 0.4673129117 | 0.8678849970 |
| 51 | -0.6358819240 | -0.2889409846 | 0.3766254117 | -0.7394900030 |
| 52 | 0.2584930760 | 0.0163715154 | -0.2472495883 | -0.5719900030 |
| 53 | -0.0421944240 | -0.7845659846 | 0.9648129117 | 0.2585724970 |
| 54 | 0.0863055760 | 0.0184340154 | -0.0138120883 | 0.3085724970 |
| 55 | 0.0091180760 | 0.0313715154 | -0.0339370883 | -0.2651775030 |

| | | | | |
|---|---|---|---|---|
| 56 | 0.1598680760 | 0.1334965154 | 0.0074379117 | 0.0691349970 |
| 57 | -0.1311319240 | 0.0387465154 | 0.2155004117 | -0.0087400030 |
| 58 | -0.3813819240 | -0.0492534846 | -0.2754370883 | 0.0946349970 |
| 59 | 0.3343680760 | -0.0148784846 | -0.0678120883 | 0.0819474970 |
| 60 | -0.0251319240 | -0.0315034846 | -0.0011870883 | 0.0270099970 |

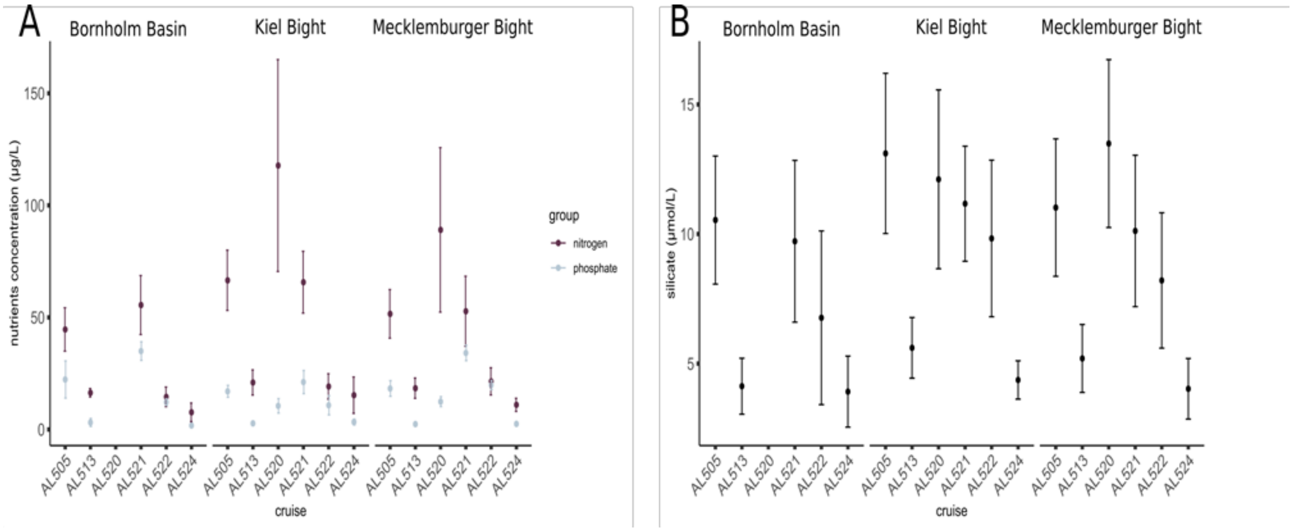


**Fig S2.** Nutrient concentrations at sampling locations during the cruises. (A) Total nitrogen (nitrate and nitrite), in purple and phosphate, in grey. Concentrations are expressed in µg $L^{-1}$ while silicates (B) are expressed in µmol $L^{-1}$. Kiel Area is hereby presented as Mecklenburger Bight and Kiel Basin separately, to highlight the overall non-significant differences between the two areas (hence, throughout considered unitary as Kiel Area). Error bars indicate ±SD around the mean.

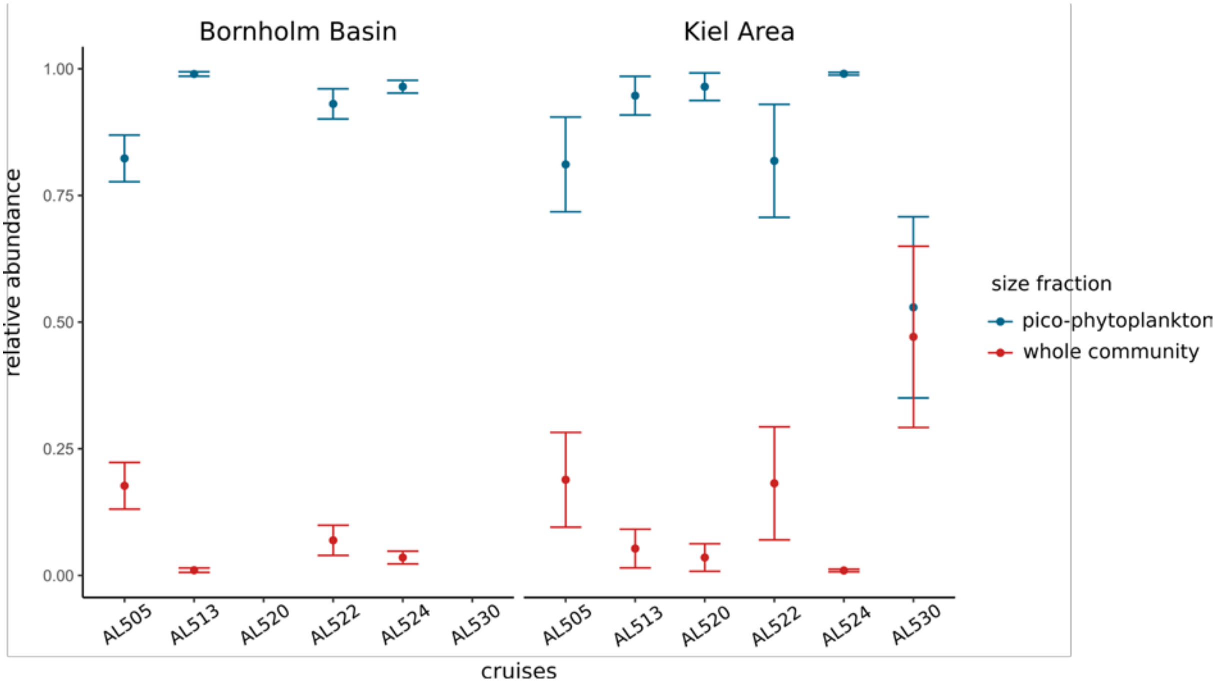


**Fig S3**. Relative abundance of pico-phytoplankton (in blue) and bigger (> 3 µm in diameter) organisms (in red) composing the whole community throughout the cruises. Contribution of the bigger cells was calculated from cytometer enumeration of events bigger than reference size beads. Error bars indicate ±SD around the mean.

**Table S4.** Outcome of model selections. Mean sampling temperature (meanT), size fraction (fraction), geographical areas (geo) and interaction between size fraction and geographical areas (fraction:geo) are here considered as fixed effects. The first line corresponds to the most parsimonious model. A plus states the variable is even slightly significant. Reduction was carried out using the function *dredge* in the *MuMIn* package (1.43.6) in the R environment. Models are ranked according to the AICc scores. Where delta AICc <2, models are averaged until delta AICc exceeds 2. (A,C) were fitted on the thermal optima for gross photosynthesis, excluding and including the heat wave event respectively. (B,D) were fitted on thermal optima for respiration, excluding and including the heatwave event respectively. (df: degree of freedom; logLik: log-likelihood; delta: delta AIC). The averaged models are highlighted in bold.

**A** Thermal optima gross photosynthesis excluding heatwave

| Intercept | fraction | geo | meanT | fraction:geo | df | logLik | AICc | delta | weight |
|---|---|---|---|---|---|---|---|---|---|
| **8.43** | **+** | **NA** | **0.61** | **NA** | **5.00** | **-125.24** | **262.30** | **0.00** | **0.56** |
| 10.21 | + | + | 0.55 | NA | 6.00 | -124.88 | 264.39 | 2.09 | 0.20 |
| 12.93 | NA | NA | 0.41 | NA | 4.00 | -129.02 | 267.21 | 4.92 | 0.05 |
| 10.47 | + | + | 0.55 | + | 7.00 | -124.84 | 267.29 | 4.99 | 0.05 |
| 14.42 | + | NA | NA | NA | 4.00 | -129.12 | 267.41 | 5.12 | 0.04 |
| 16.81 | + | + | NA | NA | 5.00 | -127.83 | 267.47 | 5.17 | 0.04 |
| 16.49 | NA | NA | NA | NA | 3.00 | -131.00 | 268.68 | 6.38 | 0.02 |
| 13.82 | NA | + | 0.38 | NA | 5.00 | -128.94 | 269.70 | 7.41 | 0.01 |
| 18.17 | NA | + | NA | NA | 4.00 | -130.43 | 270.03 | 7.74 | 0.01 |
| 16.83 | + | + | NA | + | 6.00 | -127.82 | 270.27 | 7.98 | 0.01 |

**B** Thermal optima respiration excluding heatwave

| Intercept | fraction | geo | meanT | fraction:geo | df | logLik | AICc | delta | weight |
|---|---|---|---|---|---|---|---|---|---|
| **15.72** | **NA** | **NA** | **NA** | **NA** | **3** | **-119.27** | **245.26** | **0.00** | **0.42** |
| **14.42** | **NA** | **NA** | **0.17** | **NA** | **4** | **-118.90** | **247.05** | **1.79** | **0.17** |
| 15.31 | NA | + | NA | NA | 4 | -119.21 | 247.68 | 2.42 | 0.13 |
| 15.94 | + | NA | NA | NA | 4 | -119.24 | 247.73 | 2.47 | 0.12 |
| 13.92 | NA | + | 0.17 | NA | 5 | -118.83 | 249.60 | 4.34 | 0.05 |
| 14.61 | + | NA | 0.16 | NA | 5 | -118.88 | 249.70 | 4.44 | 0.05 |
| 15.53 | + | + | NA | NA | 5 | -119.18 | 250.30 | 5.04 | 0.03 |
| 14.12 | + | + | 0.17 | NA | 6 | -118.81 | 252.41 | 7.16 | 0.01 |
| 15.17 | + | + | NA | + | 6 | -119.13 | 253.06 | 7.80 | 0.01 |
| 13.86 | + | + | 0.16 | + | 7 | -118.77 | 255.41 | 10.15 | 0.00 |

**C** Thermal optima gross photosynthesis including heatwave

| Intercept | fraction | geo | meanT | fraction:geo | df | logLik | AICc | delta | weight |
|---|---|---|---|---|---|---|---|---|---|
| **9.06** | **+** | **NA** | **0.58** | **NA** | **5** | **-166.43** | **344.22** | **0.00** | **0.40** |
| **11.26** | **+** | **+** | **0.56** | **+** | **7** | **-163.82** | **344.31** | **0.10** | **0.38** |
| 9.53 | + | + | 0.56 | NA | 6 | -166.38 | 346.72 | 2.50 | 0.11 |
| 12.37 | NA | NA | 0.49 | NA | 4 | -169.29 | 347.46 | 3.25 | 0.08 |
| 11.91 | NA | + | 0.50 | NA | 5 | -169.26 | 349.88 | 5.67 | 0.02 |
| 19.50 | + | + | NA | + | 6 | -170.24 | 354.43 | 10.22 | 0.00 |
| 17.97 | NA | NA | NA | NA | 3 | -174.05 | 354.61 | 10.40 | 0.00 |
| 16.54 | + | NA | NA | NA | 4 | -173.02 | 354.93 | 10.72 | 0.00 |
| 17.99 | + | + | NA | NA | 5 | -172.21 | 355.78 | 11.56 | 0.00 |
| 19.12 | NA | + | NA | NA | 4 | -173.70 | 356.30 | 12.08 | 0.00 |

**D** Thermal optima respiration including heatwave

| Intercept | fraction | geo | meanT | fraction:geo | df | logLik | AICc | delta | weight |
|---|---|---|---|---|---|---|---|---|---|
| **10.77** | **NA** | **NA** | **0.76** | **NA** | **4** | **-164.01** | **336.92** | **0.00** | **0.55** |
| 9.86 | NA | + | 0.77 | NA | 5 | -163.79 | 338.98 | 2.06 | 0.20 |
| 10.41 | + | NA | 0.76 | NA | 5 | -163.92 | 339.24 | 2.32 | 0.17 |
| 9.56 | + | + | 0.77 | NA | 6 | -163.72 | 341.45 | 4.53 | 0.06 |
| 9.77 | + | + | 0.77 | + | 7 | -163.70 | 344.13 | 7.20 | 0.02 |
| 19.02 | NA | NA | NA | NA | 3 | -173.86 | 354.25 | 17.33 | 0.00 |
| 18.49 | + | NA | NA | NA | 4 | -173.73 | 356.37 | 19.44 | 0.00 |
| 18.97 | NA | + | NA | NA | 4 | -173.86 | 356.62 | 19.70 | 0.00 |
| 18.49 | + | + | NA | NA | 5 | -173.73 | 358.85 | 21.93 | 0.00 |
| 18.40 | + | + | NA | + | 6 | -173.73 | 361.45 | 24.53 | 0.00 |

**Table S5.** PerMANOVA results based on Bray-Curtis dissimilarities using cytometric fingerprints data in relation to geographical area of origin and size fractions including heatwave conditions (A) and excluding them (B). (Df: degree of freedom; SumsOfSqs: sum of squares; MeanSqs: mean squares)

### **A** including heatwave

| **Kiel Area pico-phytoplankton** | | | | | | |
|---|---|---|---|---|---|---|
| | **Df** | **SumsOfSqs** | **MeanSqs** | **F.Model** | **R2** | **Pr(>F)** |
| treat_cruise | 5.00 | 43105000000.00 | 8620959637 | 3.33 | 0.37 | 0.026 * |
| Residuals | 28.00 | 72591000000.00 | 2592528026 | | 0.63 | |
| Total | 33.00 | 115700000000.00 | | | 1.00 | |
| **Kiel Area whole community** | | | | | | |
| | **Df** | **SumsOfSqs** | **MeanSqs** | **F.Model** | **R2** | **Pr(>F)** |
| treat_cruise | 5.00 | 1.2256E+13 | 2.4513E+12 | 1.38 | 0.19 | 0.17 |
| Residuals | 29.00 | 5.1425E+13 | 1.7733E+12 | | 0.81 | |
| Total | 34 | 6.3682E+13 | | | 1.00 | |
| **Bornholm Basin pico-phytoplankton** | | | | | | |
| | **Df** | **SumsOfSqs** | **MeanSqs** | **F.Model** | **R2** | **Pr(>F)** |
| treat_cruise | 3 | 2.18E+10 | 7273632458 | 2.6005 | 0.31456 | 0.001 *** |
| Residuals | 17 | 4.75E+10 | 2796972802 | | 0.68544 | |
| Total | 20 | 6.94E+10 | | | 1 | |
| **Bornholm Basin whole community** | | | | | | |
| | **Df** | **SumsOfSqs** | **MeanSqs** | **F.Model** | **R2** | **Pr(>F)** |
| treat_cruise | 3 | 8.24E+09 | 2746347626 | 4.9674 | 0.46712 | 0.001 *** |
| Residuals | 17 | 9.40E+09 | 552877173 | | 0.53288 | |
| Total | 20 | 1.76E+10 | | | 1 | |

### **B** excluding heatwave

| **Kiel Area pico-phytoplankton** | | | | | | |
|---|---|---|---|---|---|---|
| | **Df** | **SumsOfSqs** | **MeanSqs** | **F.Model** | **R2** | **Pr(>F)** |
| treat_cruise | 4 | 3.33E+10 | 8313790358 | 2.6376 | 0.31447 | 0.056 . |
| Residuals | 23 | 7.25E+10 | 3151987654 | | 0.68553 | |
| Total | 27 | 1.06E+11 | | | 1 | |
| **Kiel Area whole community** | | | | | | |
| | **Df** | **SumsOfSqs** | **MeanSqs** | **F.Model** | **R2** | **Pr(>F)** |
| treat_cruise | 4 | 1.18E+13 | 2.9511E+12 | 1.3773 | 0.18669 | 0.201 |
| Residuals | 24 | 5.14E+13 | 2.1427E+12 | | 0.81331 | |
| Total | 28 | 6.32E+13 | | | 1 | |
| **Bornholm Basin pico-phytoplankton** | | | | | | |
| | **Df** | **SumsOfSqs** | **MeanSqs** | **F.Model** | **R2** | **Pr(>F)** |
| treat_cruise | 2 | 1.85E+10 | 9264437717 | 2.533 | 0.28042 | 0.008 ** |
| Residuals | 13 | 4.75E+10 | 3657429550 | | 0.71958 | |
| Total | 15 | 6.61E+10 | | | 1 | |
| **Bornholm Basin whole community** | | | | | | |
| | **Df** | **SumsOfSqs** | **MeanSqs** | **F.Model** | **R2** | **Pr(>F)** |
| treat_cruise | 2 | 6.41E+09 | 3206027949 | 4.4379 | 0.40573 | 0.001 *** |
| Residuals | 13 | 9.39E+09 | 722423819 | | 0.59427 | |
| Total | 15 | 1.58E+10 | | | 1 | |
